# Estimates of the rate of adaptive evolution in four wild populations of *Nemophila menziesii*

**DOI:** 10.64898/2026.09.21.753227

**Authors:** Helen E. Payne, Devin E. Gamble, Lisa Kim, Anna R. H. Peschel, Susan J. Mazer

## Abstract

Populations that have detectable additive genetic variance for fitness (V_A_(W)) have the capacity for increase in their mean fitness through natural selection. The extent to which the magnitude of V_A_(W) in wild populations is predictive of the rate of adaptation is not well known, because few empirical field-based studies have been designed to address this question. Here, we report a two-year field experiment with pedigreed seeds embedded in four natural populations of the annual wildflower Baby Blue Eyes (*Nemophila menziesii* H. & A.), a species native to western North America. We first estimated V_A_(W) in a parental generation and then evaluated whether that estimate accurately predicted the adaptive response observed in the subsequent offspring generation grown under natural field conditions. We achieved this by directly comparing the mean fitnesses of the parental generation and their progeny grown under the same environmental conditions. We detected V_A_(W) and evidence of adaptation in all populations despite inter-annual differences in environmental conditions that directly affected fitness. Genotype-by-year interactions in fitness indicate that the fitness ranking of families differed between years, implying that the selective environment shifted over time. Additionally, although the environment in the second year reduced population mean fitness in three sites, the offspring generation maintained mean fitness at or above replacement, suggesting that adaptive evolution may have buffered the effects of environmental change. Overall, our results suggest that the four populations harbored sufficient V_A_(W) for adaptation through natural selection to the environmental conditions observed during the two years of this study.

**Teaser Text:** When does adaptive potential lead to adaptation? In this study of Baby Blue Eyes (*Nemophila menziesii)*, we planted seeds of known parentage from four natural populations into their site of origin and, in the following year, sowed the same sibships alongside their offspring. We evaluated the potential for adaptation in each population and the extent to which it was realized in the offspring generation. For all populations, we detected a significant capacity for adaptive evolution, which was expressed in the second generation. In all populations, the combination of survivorship and offspring production was sufficient for population replacement (i.e., mean fitness ≥ 1). Genotype-by-year interactions among sire effects on fitness suggest that the selective environment differed between years. Our study provides evidence that wild populations in their home environment possess the capacity for adaptation due to natural selection over short time scales.

## Introduction

Additive genetic variance (V_A_) quantifies the allelic variation in a quantitative trait that is available to support response to selection (Falconer & Mackay, 1996). Fitness (*W*), characterized here as lifetime female fecundity (total seed production) and survival, is the phenotype through which natural selection operates. Consequently, a population’s additive genetic variance for fitness (V_A_(W)) is proportional to a population’s capacity for adaptive evolution, which is a change in mean fitness between subsequent generations resulting from natural selection (Fisher, 1930; Houle, 1992, 1998; Lande & Shannon, 1996) Populations with high V_A_(W) are expected to have a greater capacity for adaptation than those with low V_A_(W) (Arnold, 2023).

V_A_(W) is challenging to quantify in field settings, but existing studies indicate that V_A_(W) can be substantial in wild populations. Field studies of the annual plant *Chamaecrista fasciculata* (Michx.) Greene (Fabaceae) have detected substantial V_A_(W) for populations growing in manipulated environments (Peschel et al., 2021), in novel sites (Peschel & Shaw, 2024; Sheth et al., 2018), and in their home environment (Kulbaba et al., 2019; Shaw et al., 2026). Additionally, Bonnet et al. (2022) analyzed long-term lifetime breeding success data from 19 pedigreed wild animal populations and found that often populations harbor significant additive genetic variance in fitness, suggesting that rapid adaptive evolution can occur under natural conditions.

Estimating both V_A_(W) and mean fitness (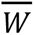) is necessary to evaluate not only whether populations possess the capacity to adapt, but also whether adaptive responses are sufficient to maintain population growth under changing environmental conditions. For populations to persist, their growth rate must, on average, be at or above replacement (i.e., mean fitness > 1) even in the absence of detectable adaptation if existing genotypes are sufficiently tolerant of prevailing environmental conditions or buffer environmental variation through plastic responses (Cross & Eckert, 2024). Alternatively, V_A_(W) may be present but not sufficient to support adaptive response that maintains a positive population growth rate in the subsequent generation.

The presence of additive genetic variance in fitness (V_A_(W)) alone does not guarantee that adaptive evolution will occur. Although (V_A_(W)) determines a population’s capacity to respond to natural selection, a genetically based increase in mean fitness (i.e., adaptation) will only be realized if the selective environment remains sufficiently consistent for fitness differences among genotypes to translate into evolutionary change. Changes in mean fitness can also arise solely from environmental variation and therefore should not be interpreted as evidence of adaptation. Temporal environmental variation can alter both the strength and direction of selection among genotypes, such that genotypes favored in one generation have reduced fitness in the next, thereby weakening or preventing the expected adaptive response. Our understanding of the degree to which the magnitude of V_A_(W) is predictive of the rate of adaptation is limited because there are only two published studies that have directly compared estimates of V_A_(W) to the change in mean fitness between generations. Peschel & Shaw (2024) and Shaw et al., (2026) evaluated wild populations of the annual legume *Chamaecrista fasciculata*, and found the magnitude of V_A_(W) to poorly predict the mean fitness of the progeny generation. Even though the environment was not experimentally manipulated between generations, both studies detected evidence for Genotype × Year (G × Y) interactions, indicating that the selective environment changed between generations. They argue that the change in environment was likely partly responsible for deviations between the predicted and the realized rate of adaptation.

Here, we build upon Peschel and Shaw (2024) and Shaw et al. (2026) by evaluating both the capacity for adaptive evolution and the extent to which adaptation is realized by conducting a two-generation field experiment in which parental and offspring generations of the annual herb, *Nemophila menziesii* H. and A. (Hydrophyllaceae) were grown side by side in four wild populations. This design provides a rare opportunity to assess whether populations with greater additive genetic variance in fitness exhibit greater realized adaptive responses between generations. Our focal populations span a latitudinal precipitation gradient in California, USA, providing insight into variation in adaptive capacity among populations of the same species from naturally differing environmental regimes. We addressed the following questions:*(1)* Is V_A_(W) detectable in each of the four natural populations? *(2)* How does the estimated magnitude of V_A_(W) in a given population compare to the genetic change in 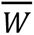 between generations? *(3)* Are populations tolerant of the environment such that they persist at or above replacement under stressful conditions, and/or does adaptive evolution increase population growth rates?? *(4)* Within populations, is there evidence that Genotype × Year interactions constrain the response to selection?

## Methods

### Study Organism

*N. menziesii* is native to North America and occurs along the west coast of Oregon and California (Cruden, 1974; McCall & Barr, 2012) in organic, well-drained soils at elevations below 6,500 feet (Cruden, 1974). Flowering phenology in *N. menziesii* varies among populations and across latitudinal and elevational gradients, with higher-elevation populations generally flowering later and for longer durations than lower-elevation populations (Cruden, 1972). *N. menziesii* includes several recognized varieties (*var. menziesii, var. atomaria, and var. integrifolia*), which differ in floral morphology, particularly petal color and spotting (Patterson & Halse, 2021). All individuals in this study were identified broadly as *N. menziesii*, and no formal differentiation by variety was made during seed collection or experimental design.

The majority of *N. menziesii* individuals are hermaphroditic and produce large, protandrous flowers, in which pollen is released before the stigma becomes receptive, that are outcrossed by a wide range of pollinating insects, including bees and flies (Andersson, 1994; Cruden, 1972). Although self-pollination can occur, the vast majority of greenhouse-raised plants do not produce seed in the absence of pollinators (Gomez & Shaw, 2006). Most individuals bear bisexual flowers, but some are strictly female, bearing male-sterile flowers that do not produce pollen (Gomez & Shaw, 2006), and many individuals produce some male-sterile flowers under greenhouse conditions (SJM, personal observation).

In our study sites (Figure 1), we observed *N. menziesii* seeds germinating in November, with flowering starting in February and continuing through June. Fruit capsules matured and dehisced from March to June (*Nemophila menziesii*, 2023). Seeds may remain dormant in the seed bank for several years until local temperature and precipitation conditions trigger germination (Cruden, 1972).

**Figure 1.**
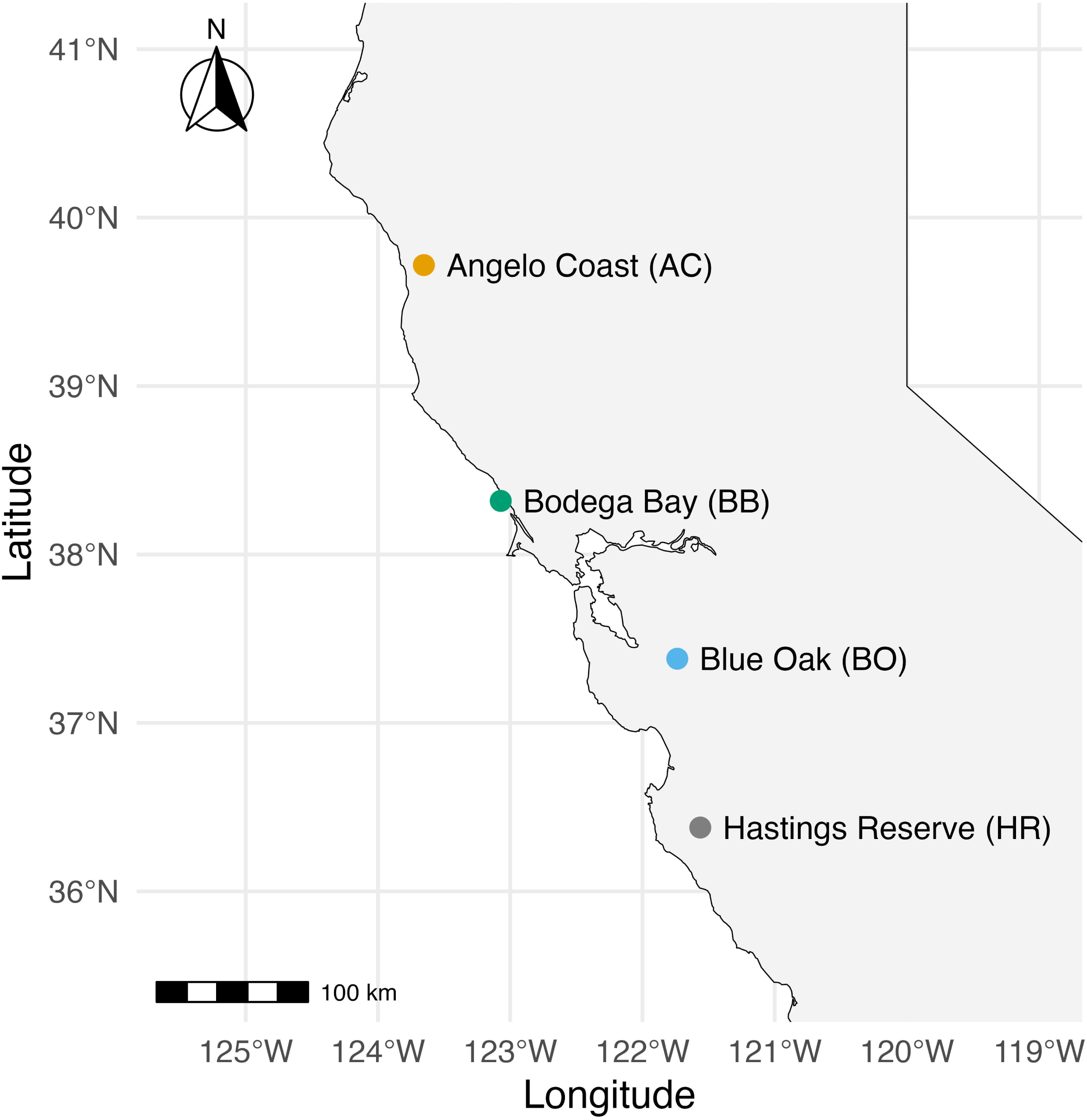
Locations of the field sites used in this study were Angelo Coast Range Reserve (Angelo Coast, AC), Bodega Bay Marine Lab (Bodega Bay, BB), Blue Oak Ranch Reserve (Blue Oak, BO), and Hastings Natural History Reservation (Hastings Reserve, HR). The Angelo Coast Range Reserve is ∼500 kilometers north of the southernmost site, Hastings Natural History Reservation. See Table 1 for latitude and longitude.

### Study populations

The focal populations for this study were located at four University of California Natural Reserve System (UCNRS) reserves where *N. menziesii* naturally occurs.

These sites were chosen along a north-south gradient (Figure 1), resulting in climatic differences among them in seasonal and annual means (Table 1). Although climate was the primary consideration in site selection, other environmental factors also vary among sites and may contribute to differences among populations. Real-time environmental sensors were present at each field station, and data were archived in the historical Dendra records (Dendra Science, 2026).

**Table 1.** GPS coordinates and climatic data for each study site: Angelo Coast (AC), Bodega Bay (BB), Blue Oak (BO), and Hastings Reserve (HR). Mean annual and spring temperature in degrees Celsius, cumulative annual and spring precipitation (PPT) in millimeters (mm), latitude, longitude, and elevation (meters). Mean spring temperature was calculated as the average of mean daily temperatures for March, April, and May over a 24-hour period. Climatic data are based on 30-year climate normals (means) (± SD) observed from 1991 to 2020 obtained from ClimateNA (Wang et al., 2016).

| Site | Latitude | Longitude | Annual PPT(mm) | Spring PPT (mm) | Annual Temp (°C) | Spring Temp (°C) | Elevation (m) |
| --- | --- | --- | --- | --- | --- | --- | --- |
| AC | 39.7469 | -123.6380 | 2135.6 ± 637.1 | 541.4 ± 255.8 | 12.3 ± 0.58 | 10.5 ± 1.1 | 567 |
| BB | 38.3146 | -123.0694 | 910.5 ± 262.9 | 222.7 ± 118.6 | 12.43 ± 0.53 | 11.7 ± 0.9 | 2 |
| BO | 37.3870 | -121.7158 | 670.4 ± 203.5 | 185.0 ± 101.7 | 14.77 ± 0.58 | 12.4 ± 1.1 | 728 |
| HR | 36.378<br>3 | -121.566<br>2 | 729.9 ±<br>222.5 | 187.5 ±<br>100.9 | 14.78 ±<br>0.59 | 12.4 ±<br>1.0 | 603 |

### Seed sampling to generate pedigreed seed

At each site, seeds were collected from a minimum of 200 maternal plants in spring 2018. To reduce the probability that these maternal plants were closely related, seeds were collected from individual plants growing at least one meter apart. The seeds of each maternal plant were collected separately in paper coin envelopes.

### Breeding design to produce pedigree populations in the greenhouse

From August 2018 to July 2019, one offspring from each field-sampled maternal plant was reared in the greenhouse and used in hand-pollinations to generate a pedigreed population of seeds (see S1 for details of plant cultivation and greenhouse conditions). We used a paternal half-sibling design (details below), which enables estimation of the variance component due to the effect of sires (male parents) on progeny fitness, with the expectation that this variance component represents one-quarter of V_A_(W) (Falconer & Mackay, 1996). The greenhouse was not large enough to simultaneously accommodate all individuals to be pollinated, so we raised and pollinated a subset of the field-collected maternal families from each population in seven successive cohorts from August 2018 to December 2019 (see supplementary material for details). In each cohort, from each population, we cultivated two seeds representing each of approximately 40 field-collected maternal seed families. Upon flowering, these plants were assigned at random to individual mating groups, each composed of one pollen donor and three pollen recipients derived from different field-collected maternal families. To the extent possible, only one seed from each field-collected maternal family was used. However, to ensure that each mating group included co-flowering individuals, in some cases, the two seeds cultivated from a maternal family were required to be used. In such cases, one of the two offspring was used as a pollen donor in one mating group, and the other was used as a pollen recipient in a different mating group. The number of maternal seed families used for both donor and recipient roles (i.e., two) differed among populations (AC: 18, BB: 3, BO: 21, HR: 34). To preserve pedigree independence and minimize inbreeding, no field-collected maternal family was used more than once as a pollen donor or as both a pollen donor and recipient within the same mating group. In each population, across all cohorts, ∼50 individuals were randomly designated as pollen donors (sires) and 150 as pollen recipients (dams).

The pollen from each pollen donor was used to hand-pollinate ∼100 flowers on each of three maternal plants (Figure 2). Throughout the flowering period of each pollen recipient, the petioles of its unopened flower buds were labeled with narrow strips of tape to indicate that they were to be pollinated. Then, using needle-nose forceps, all buds were gently pried open and emasculated (i.e., the stamens were removed from the buds before they naturally opened) to ensure that subsequently collected fruits arose from deliberate pollinations. Flowers that had not been emasculated were removed.

**Figure 2.**
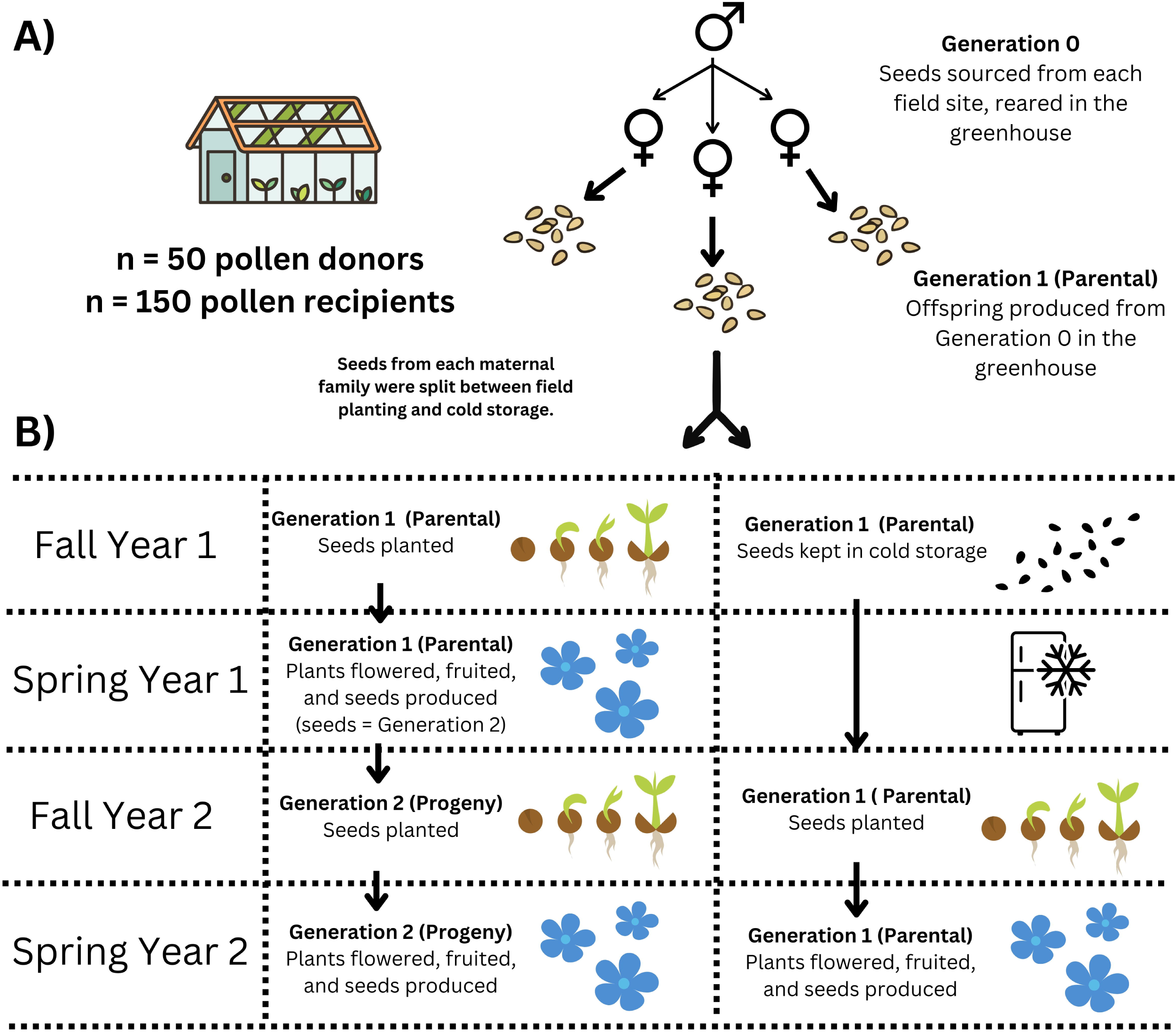
Experimental design for estimating V_A_(W) and evaluating the realized rate of adaptation at each site. A) Field-collected seeds of *N. menziesii* plants were reared in the greenhouse and, when flowering, assigned as either pollen donors (sires, paternal plants) or pollen recipients (dams, maternal plants). Each pollen donor was randomly assigned to pollinate three distinct maternal plants. This nested breeding design produced ∼40-50 paternal sibships and ∼107-150 maternal sibships (all seeds produced by a single maternal plant) per population, which comprise generation 1. Each pollen donor was used to hand-pollinate up to 100 flowers on each of the three dams nested within it. B) Greenhouse-produced seeds (generation 1) were divided into two groups: one group was planted in our field sites in Fall 2021 and monitored in Spring 2022, while the other remained in cold storage until Fall 2022. Seeds produced by generation 1 plants were designated as generation 2 (the progeny generation) and planted in Fall 2022 simultaneously with the reserved generation 1 seeds. This design allowed direct comparison of generation 1 and 2 individuals growing under the same conditions in Fall 2022-Spring 2023. This design was applied to all four populations.

As controls to assess unintended pollination in the greenhouse, 13 plants from Cohort 4 were used, comprising three or four plants per population. All flowers on these plants were emasculated during development, and the plants were placed throughout the greenhouse among the experimental pollen recipients. Fruit production on these 13 embedded plants was assessed, and the near-zero fruit set confirmed that unintended pollination was extremely limited. The 13 plants produced a total of 1,651 flowers, of which 0.02% set fruit; greenhouse conditions are described in the supplemental material (Table S1.1). Moreover, given that we hand-pollinated each emasculated bud within a few hours of stigma receptivity, the opportunity for insect pollination was very low because compatible pollen had already been deposited on the stigmas.

As fruits of the hand-pollinated maternal plants ripened, the seeds from each maternal family were collected and stored in coin envelopes separately, and placed with silica gel in zip-lock plastic bags. These pedigreed seeds were placed in cold storage (9°C) until fall 2021, when the first year of the field experiment was initiated. Pedigreed seeds produced from the greenhouse-reared plants are referred to as generation 1.

### Experimental field planting design — Year 1

In Fall 2021 (hereafter referred to as “year 1”), a portion of the generation 1 seeds representing each population was used as follows (see Figure S7 for a visual representation of the plot set up): At each site, three blocks were configured, each containing 3-4 transects along the length of which 1-meter linear segments were established. At AC, 9 transects were established, each with ∼36 segments; at BO, 12 transects were established, each with ∼23-44 segments; and at HR, 9 transects were established, each with ∼46 segments. To minimize confusion between pedigreed individuals and naturally occurring plants from the source population, and to facilitate locating experimental individuals during surveys, surface vegetation and litter were cleared for 5 cm on both sides along the length of a given 1-meter segment (creating a cleared 10 cm x 1-meter area). Each maternal sibship was randomly assigned to a 1-meter segment within each block. Along the length of each segment, 8 seeds (10 seeds at AC) from the assigned maternal plant were planted 10 cm apart. To ensure accurate identification and tracking of planted seeds, we planted individual seeds into separate peat and coconut fiber plugs inserted into the soil and marked each planting location with labeled skewers (see supplemental material for details on the seed planting protocol). This approach minimized confusion of our experimental seeds with those from the seed bank and reduced seed displacement due to rainfall, improving our ability to locate and monitor the pedigreed seeds.

Each segment was spaced approximately one meter from neighboring segments, and transects were placed ∼five meters apart to distinguish each transect and facilitate access by providing space for observers. At BB, the patchy distribution of suitable habitat for *N. menziesii* precluded the use of transects. Instead, we identified and georeferenced 156 locations within 1.5 meters of individuals of the small shrubs, *Eriogonum latifolium* and/or *Lupinus arboreus,* which appear consistently to serve as nurse plants for *N. menziesii* at BB, and were used to identify suitable sites for establishment. At each of these locations, one to three 1-meter segments were established, with each segment containing the seeds of one maternal sibship. We sowed 3210 seeds at AC, 3528 seeds at BB, 3168 seeds at BO, and 3288 seeds at HR.

### Experimental field planting design — Year 2

In Fall 2022 (hereafter referred to as “year 2”), both the reserved generation 1 seeds (those in cold storage) and the open-pollinated seeds collected from the field-grown plants sown in year 1 (i.e., generation 2 seeds) were sown at each field site using the same spatial design as in year 1 (Figure 2, Figure S7). The transects and one-meter segments established in year 1 were reused in year 2. However, in year 2, each segment contained two maternal sibships of five seeds each, rather than a single maternal sibship of eight seeds (10 seeds at AC). All sibships from generation 1 and generation 2 were randomized to ensure that generation 1 and generation 2 sibships were interspersed throughout the transects rather than clustered by generation or family. Labeled bamboo skewers were placed beside each seed at planting to distinguish individuals. We sowed 3195 generation 1 and 2 seeds at AC, 2205 seeds at BB, 3090 seeds at BO, and 4055 seeds at HR.

### Field-collected data to estimate mean lifetime fitness

At each site, we estimated mean lifetime fitness for every pedigreed generation in each year. For each seed planted, we recorded whether it germinated and survived to flower, after which we continued monitoring a subset of individuals by randomly selecting one individual that survived to flowering from each segment (hereafter referred to as the fitness plant). For these individuals, we recorded total lifetime fruit production and total lifetime seed production (Figure S8).

A plant was considered to have survived to flowering if it was in flower or bore pedicels indicating previous flowering. Fruits of *N. menziesii* dehisce upon reaching maturity; therefore, fruits from fitness plants were collected prior to dehiscence whenever possible and classified as intact. Fruits that had dehisced or showed evidence of herbivory were classified as open. Total fruit production was defined as the sum of the numbers of open and intact fruits produced over a plant’s lifetime. For all intact fruits, we counted the number of filled seeds they contained.

*Statistical Analysis:*

All analyses were performed in R version 4.2.3 (R Core Team, 2023). Aster analysis was conducted using the R package “aster” (version 1.1-3, Geyer, 2025).

### Aster models to estimate lifetime fitness and additive genetic variance for fitness

Lifetime fitness is challenging to model accurately because its frequency distribution within populations does not conform to known statistical distributions. Many individuals die or produce no seeds, resulting in zero inflation, and fitness distributions are often multimodal. Aster models validly estimate lifetime fitness and its standard error by modeling fitness components that follow known statistical distributions, and jointly analyzing them while taking into consideration the dependence of successive life-history stages on those prior (Geyer, 2023; Shaw et al., 2008). The aster graphical model included the following nodes, each indicating a point during the life cycle when data were recorded or individual plants were selected for sampling, and each with a specific type of distribution: survival to flowering (Bernoulli distribution), designation of the fitness plant (Bernoulli distribution; subsampling node), total number of fruits produced by the fitness plant (Poisson distribution), number of closed fruits produced by fitness plant (Bernoulli distribution; subsampling node), and total number of seeds contained within the closed fruits of the fitness plant (Poisson distribution) (See a visual representation of the model in Figure S8).

Effects of individual pollen donors on offspring fitness were estimated separately, based on the above model, following Peschel and Shaw (2024) and Geyer (2022). Paternal effects estimate the average additive effect of a donor’s alleles across multiple dams, thereby minimizing maternal environmental influences and providing an estimate of each donor’s additive genetic contribution to offspring fitness as expressed in the study environment.

Generation 2 seeds were produced following open pollination, so the paternal parentage of the individuals in this generation was unknown, which precluded estimation of V_A_(W); therefore, the models applied to generation 2 did not include a random effect for the parent or for the maternal grandfather. 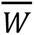 is estimated as the unconditional expected value of lifetime reproductive output for individuals in a given generation for each population, accounting for the sequential structure of the life-history transitions as well as any fixed or random effects included in the model.

For generation 1, we estimated V_A_(W) as follows. To account for any linear environment gradient within each site, we included transect as a continuous fixed effect. At BB, which lacked transects or blocks, we included location as a continuous fixed effect. Location was defined as the position of each of the 156 1-m planting segments along a single linear transect, allowing us to account for environmental variation along this spatial gradient. When modeled separately, we detected maternal and paternal random effects to be significant and of comparable magnitude.

When both were included in models, neither was significant. We combined their contributions into a single random effect, hereafter referred to as “parental variance” following Peschel and Shaw (2024) and Geyer (2022), and consistent with what was illustrated in Falconer and Mackay (1996) example 10.4.

For generation 2, we fit fixed-effect aster models to estimate 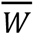, which also included transect or location as a continuous fixed effect. The realized rate of genetically based adaptation for the population at each site was estimated as the difference in 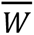 between generation 1 and 2 when both were reared in year 2 (*W_G_*_2, *Y*2_ − *W_G_*_1, *Y*2_). Because both generations were evaluated under the same environmental conditions, this comparison isolates the genetically based change in mean fitness. To quantify the environmental contribution to the change in 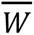 we calculated the difference in mean fitness between generation 1 grown in year 2 and generation 1 grown in year 1 (*W _G_*_1, *Y*2_− *W_G_*_1, *Y*1_). Consequently, the total difference in mean fitness between generation 1 in year 1 and generation 2 in year 2 (*W_G_*_2, *Y*2_ − *W_G_*_1, *Y*1_) reflects the combined effects of the genetic change between generations and environmental differences between years.

### Evaluation of Gene by Year (G × Y) interaction

We used three approaches to seek evidence for a genotype-by-year interaction, which gives insight into whether the genetic contributions of sire groups to V_A_(W) differed with change in environmental conditions between years. First, to formally test for a G *×* Y interaction, we compared aster models using likelihood ratio tests for each site. Both models included year and transect (for BB, location was used instead of transect) as fixed effects and a parental random effect, but the G × Y model also included the interaction between parental × year as a random effect, allowing sire effects on *W* to vary between years. The G x Y interaction was tested by a likelihood ratio test, with *p*-value <0.05.

At HR, the full 5-node random-effects model failed to converge when fit to the combined-year (year 1 + year 2) dataset and was therefore excluded from the analysis of G × Y interactions. Second, we tested for G × Y interactions in fitness by comparing models with and without a parental × year random effect. Significant interactions indicate that the genetic basis of fitness differed between years, reflecting changes in the variance among sire effects, changes in sire rankings, or both. Third, to observe and visualize the rank and the variance of the sire effects on *W* estimated in both years, we plotted estimates of the sire effects expressed by generation 1 in years 1 and 2, and calculated Pearson’s correlation coefficient for each site to assess the consistency of the sire effects on *W* between years. These correlations were used descriptively to evaluate rank consistency and do not account for uncertainty in the sire effect estimates (Peschel & Shaw, 2024).

## Results

### Environmental conditions during year 1 and year 2

There was substantially lower cumulative precipitation in year 1 than in year 2 at all sites (Table 2). Mean annual temperatures were similar between years within sites.

**Table 2.** Mean annual temperature (T) and cumulative precipitation (PPT) during year 1 and year 2, at each study site: Angelo Coast (AC), Bodega Bay (BB), Blue Oak (BO), and Hastings Reserve (HR). Data were obtained from real-time environmental sensors installed at each field station and archived in the historical Dendra records.

| Site | Cumulative annual PPT (mm) year 1 | Mean annual T (°C) year 1 | Cumulative annual PPT (mm) year 2 | Mean annual T (°C) year 2 |
| --- | --- | --- | --- | --- |
| AC | 1,410.0 | 11.0 | 1,963.2 | 11.1 |
| BB | 617.5 | 11.1 | 1,147.1 | 11.5 |
| BO | 421.4 | 13.8 | 1,010.4 | 12.7 |
| HR | 374.4 | 13.7 | 953.0 | 12.7 |

*Estimates of V_A_(W):* We detected significant V_A_(W) in all four populations of *N. menziesii* for generation 1 in both years (Table 3, column A). The magnitude of V_A_(W) for generation 1 differed between years within populations, and this difference was substantial, relative to standard errors, for all populations except AC. For BO, V_A_(W) was lower in year 2 than in year 1, while for BB and HR, V_A_(W) was higher in year 2 than in year 1. V_A_(W) also differed among populations within each year; notably, BO had V_A_(W) values that were an order of magnitude lower than those of the other populations in both years.

**Table 3.** Additive genetic variance for lifetime fitness (V_A_(W)), mean lifetime fitness (W), realized rate of adaptation, and the contributions of genetic and environmental change to generation 2 fitness in four populations of *Nemophila menziesii*. The values displayed in columns B and C are also represented in Figure 3. In column A, * indicates that V_A_(W) is significantly greater than 0, based on the significant random effect term accounting for parental variance in our random effect aster models (p-value < 0.05). Blank squares indicate values that were not estimated in year 2.

|  | A | B | C | D | E | F |
| --- | --- | --- | --- | --- | --- | --- |
| Site & year of observation | Generation 1, $V_A(W)$ (±SE) | Generation 1, $\bar{W}$ (±SE) | Generation 2, $\bar{W}$ (±SE) | Rate of adaptation | Effect of genes and environment on generation 2 $\frac{2}{\bar{W}}$ | Effect of environment on generation 2 $\frac{2}{\bar{W}}$ |
| Angelo Coast Year 1 | 5.19 (1.76)* | 14.86 (0.57) | – | 12.79 | 3.76 | -9.03 |
| Angelo Coast Year 2 | 4.37 (1.87)* | 5.83 (0.52) | 18.62 (2.40) | – | – | – |
| Bodega Bay Year 1 | 1.68 (0.37)* | 1.49 (0.19) | – | 34.48 | 48.08 | 13.60 |
| Bodega Bay Year 2 | 103.14 (22.56)*<br>† | 15.09 (1.27) | 49.57 (6.69) | – | – | – |
| Blue Oak Year 1 | 0.58 (0.18)* | 1.77 (0.16) | – | 0.44 | -0.71 | -1.15 |
| Blue Oak Year 2 | 0.15 (0.04)* | 0.62 (0.10) | 1.06 (0.28) | – | – | – |
| Hastings Reserve | 2.25 (0.68)* | 13.93 (0.40) | – | 4.40 | -3.77 | -8.17 |
| Year 1 |  |  |  |  |  |  |
| Hastings Reserve Year 2 | 4.92 (1.17)* | 5.76 (0.38) | 10.16 (1.20) | – | – | – |

*Environmental contributions to changes in W:* For all populations, the contribution of the environmental difference to the change in 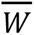 between generations exceeded the genetic contribution to change in mean fitness resulting from natural selection in the first year. This was evidenced by a greater difference in the estimates of mean fitness between the generation 1 cohorts in year 1 and 2 compared to the difference in the estimates of mean fitness between generations 1 and 2 growing in year 2 (Figure 3).

**Figure 3.**
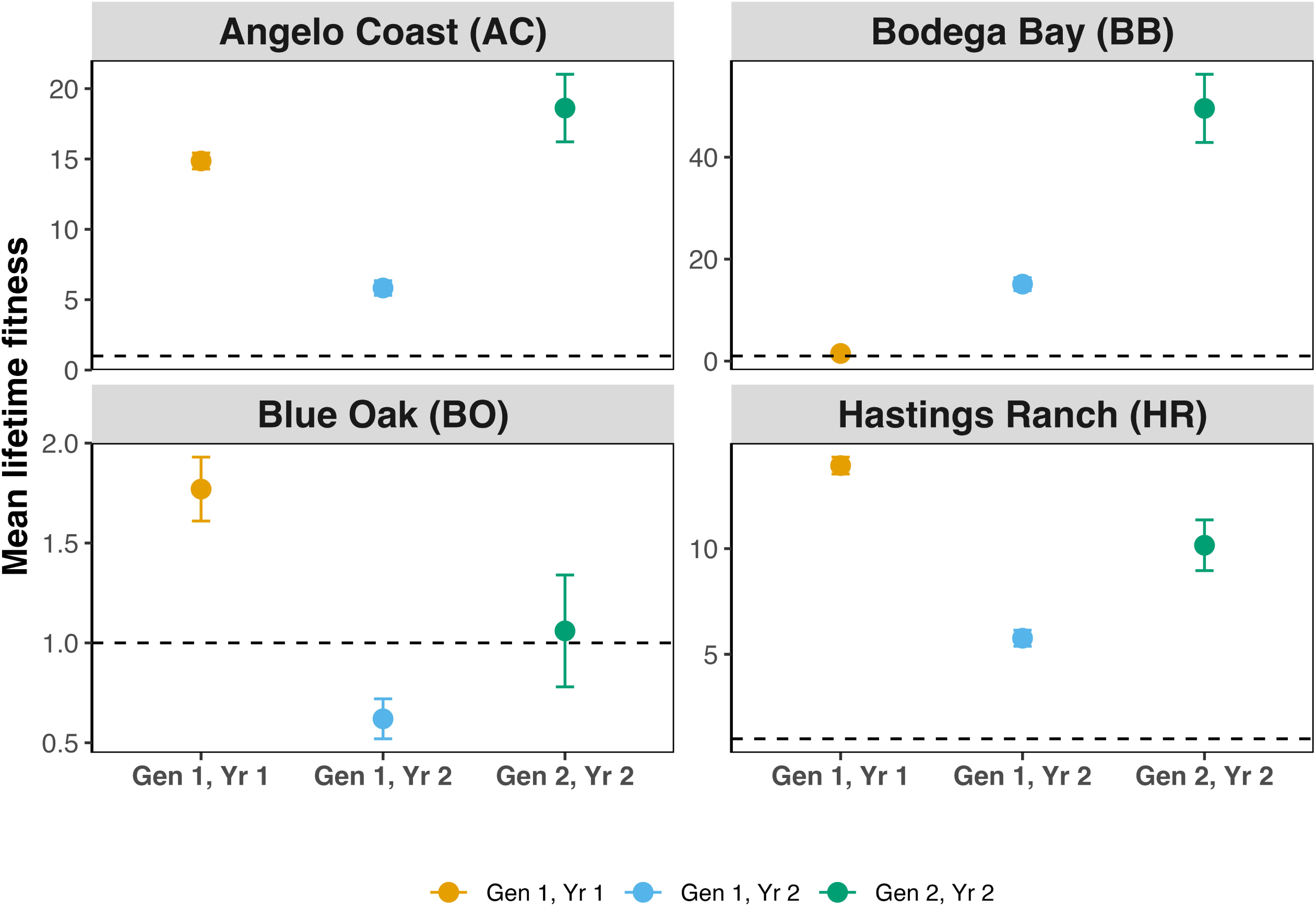
Estimates of year-specific population mean lifetime fitness (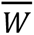) for generation 1 and generation 2 at all sites. Each panel represents one population from one site. The y-axis represents mean lifetime fitness estimated from aster models. The x-axis displays the three sources of fitness estimates: generation 1 observed in year 1 (column B from Table 3), generation 1 observed in year 2 (column B from Table 3), and generation 2 observed in year 2 (column C from Table 3). Note that the scale of the y-axis differs between populations.

Additionally, only at BB was the environment in year 2 more favorable than year 1. At this site, the estimate of 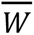 for generation 1 in year 2 was higher than the estimate in year 1 (Table 3, column F). At the remaining sites, 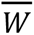 of generation 1 was lower in year 2 than in year, which suggests the environment was poorer at these sites in the second year.

### Genetic contribution to changes in W

We detected adaptation through natural selection at all sites despite the more stressful environmental conditions in the second year in three of the four sites. At all sites we detected a genetically based increase in mean fitness, indicating that selection on generation 1 in year 1 resulted in a realized fitness increase in the environment of year 2. Additionally, the fitness estimate for the progeny generation at all sites was above replacement (absolute fitness >1), though only marginally so at BO (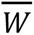 = 1.06) (Table 3, column C).

### Evaluating Genotype-by-Year (G × Y) Interactions

We detected three types of evidence for G × Y interactions. First, we directly tested for G *×* Y effects, and the model fit was significantly improved when we included a parental *×* year random effect at three sites (all p < 0.002, Table S5). The strong G x Y interactions detected at AC, BB, ,and BO indicated that the fitness of a paternal group in year 1 was not predictive of its fitness in year 2. Second, we assessed the magnitude of V_A_(W), which differed between years within populations for generation 1 at all sites, which suggests the capacity for ongoing adaptation differed between years for each population. Thirdly, we visualized these weak to negative relationships in parental breeding values observed in year 1 and year 2, which indicated that alleles associated with higher fitness in year 1 generally did not confer higher *W* in year 2 (Figure 4). At BO, we detected a negative correlation between year 1 and year 2 sire effects on mean fitness (*r* = −0.40), indicating a reversal in the relative fitness performance of sire groups across years.

**Figure 4.**
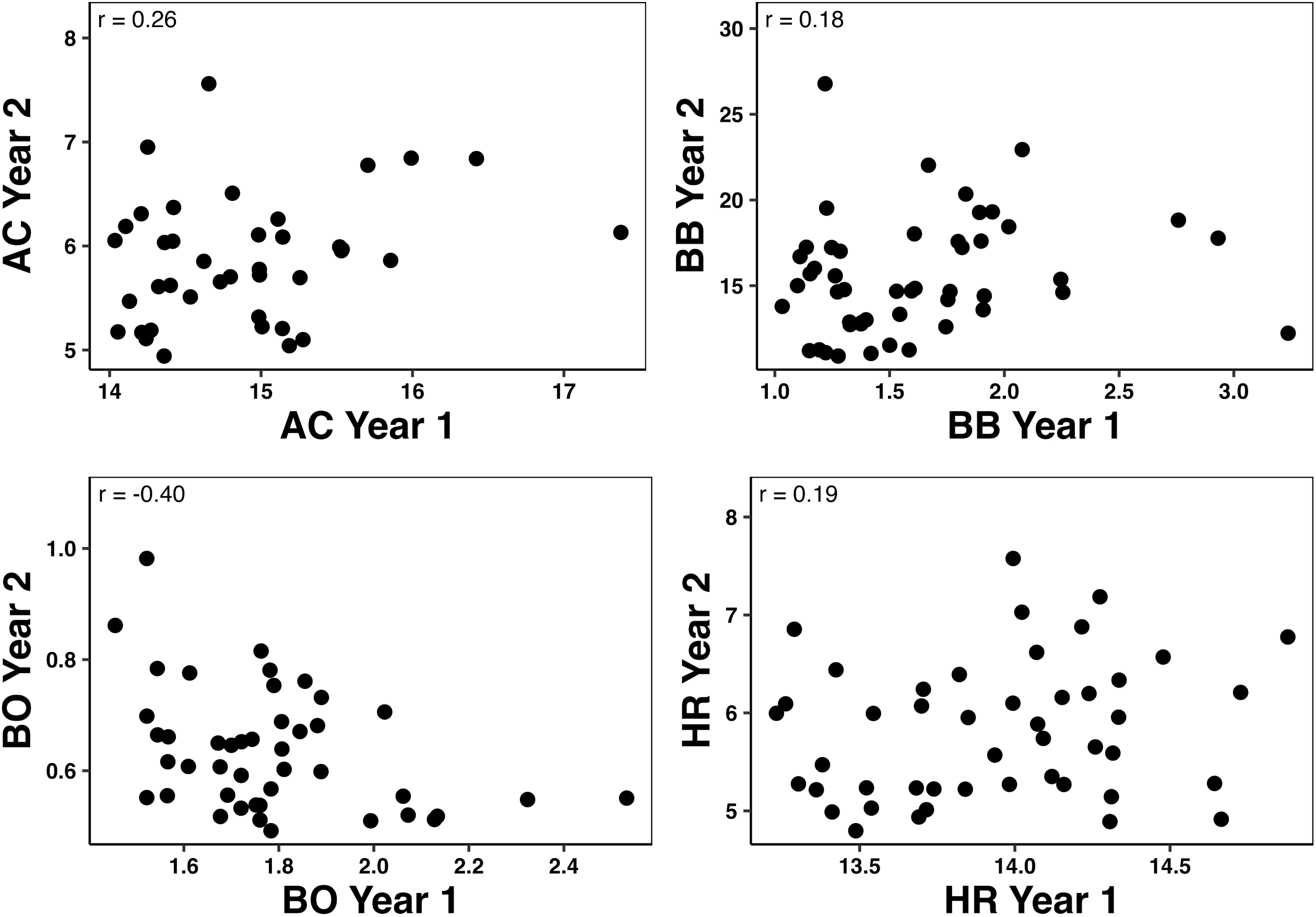
Bivariate plots of year-specific sire effects on lifetime fitness in each population. Each point represents the mean fitness of a sire group, based on the original mating group, in year 1 and year 2. Pearson’s correlation coefficients (*r*) were obtained to estimate the correlation of sire means between years, but does not include the sampling variance around the estimates of the means.

## Discussion

By quantifying additive genetic variance in fitness and mean population fitness (V_A_(W) and 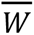) in two consecutive years for four populations of *N. menziesii*, our study provides an empirical assessment of the capacity for immediate adaptation between subsequent generations and how it varies among multiple wild populations of a single species.

### All populations had the capacity to adapt to their home environments

V_A_(W) was detectable in all four populations spanning a broad temperature and precipitation gradient, indicating that the capacity for adaptive evolution is a general feature of these populations rather than restricted to particular climate conditions; however, its magnitude differed between years within sites, indicating that this capacity is not static. This finding is notable because empirical estimates of V_A_(W) in wild populations remain relatively uncommon, particularly across environmental gradients. Our findings are consistent with the few studies reporting detectable V_A_(W) in native annual plant populations (Kulbaba et al., 2019; Peschel et al., 2021; Peschel & Shaw, 2024; Sheth et al., 2018).

### Adaptation despite environmental change between generations

All populations exhibited evidence of adaptive evolution, as the mean fitness of generation 2 exceeded that of generation 1 when grown together in year 2 environmental conditions. The extent to which adaptive responses offset environmentally induced declines in *W* varied among populations, with some populations exhibiting stronger genetically based increases in fitness than others. For example, BO showed the smallest adaptive response, consistent with its lowest V_A_(W) in both years. However, among AC, BB, and HR, the magnitude of adaptive change was not proportional to the magnitude of V_A_(W). The weak cross-year rank correlation of parental breeding values for fitness provides a likely explanation for this mismatch. That is, where the paternal families that were favored by selection in year 1 (i.e., their fitness in year 1 was relatively high) had low fitness under the environmental conditions of year 2, the realized change in mean fitness was influenced not only by V_A_(W) in year 1, but also by the change in rank of genotypes between years.

Environmental change between generations may explain deviations between the expected and realized rate of adaptation. Correlations observed between year-specific sire effects on *W* (Figure 4), reflect strong G *×* Y interactions. Thus, the alleles selected for in year 1 and transmitted to the progeny generation likely did not confer the same fitness benefit in the environment experienced by the progeny in year 2. Nevertheless, all populations exhibited evidence of adaptation, indicating that genetically based increases in *W* occurred despite shifts in the selective environment between years. However, the magnitude of these adaptive responses may have been constrained by environmentally induced declines in *W* and by G x Y interactions, which may have reduced the extent to which selection in one year translated into increased fitness in the next. These findings are similar to those of Kulbaba et al. (2019) and Peschel and Shaw (2024), who also found evidence suggesting that differences in the selective environment between years may have limited the extent to which selection in one year translates into adaptive responses in the next.

At AC, BO, and HR, changes in environmental conditions between years had a larger effect on mean fitness than the genetically based changes associated with selection. Although adaptive responses were detected, these responses were often insufficient to offset environmentally induced declines in fitness fully. Climatic conditions differed substantially between years, with generally greater precipitation in year 2 at all sites. Consistent with this shift, mean fitness, V_A_(W), and the rankings of sire breeding values differed between years, indicating that temporal environmental variation influenced both overall population performance and the genotypes favored by selection.

Differences between generations may reflect processes in addition to genetically based responses to selection, including genetic drift, gene flow, differences in seed age, maternal environments, and the open pollination that produced generation 2 (but not generation 1) seeds. Although maternal effects could contribute to generational differences, we found no evidence that maternal provisioning via seed size differed between generations: despite seed size being positively associated with survival to flowering in some populations, mean seed size did not differ significantly between generations (see Tables S6 and S7 in supplementary material). Thus, maternal provisioning through seed size is unlikely to explain the observed fitness differences.

Consequently, the realized rate of adaptation estimated here should be interpreted as the net change in mean fitness between generations grown under common environmental conditions rather than a strictly genetic response to selection.

### Population persistence despite environmental change

At three sites the environment became more stressful in the second year, with greater rainfall, as evidenced by the lower mean fitness of generation 1 in the second year compared to the first year. While environmental change between years lowered mean fitness in three of the populations, mean fitness of generation 2 remained high enough to support a positive rate of population growth (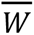 >1) for all populations.

### Stochastic Processes

Although stochastic processes such as genetic drift and gene flow can influence estimates of 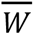, in this study, the effects of genetic drift are likely to be limited because parental individuals were randomly sampled from large, naturally occurring populations and because pedigreed populations were established using relatively large numbers of families. Under these conditions, stochastic changes in allele frequencies are less likely to overwhelm the effects of selection. All study populations were derived from the natural populations and then grown in their sites of origin and exposed to similar opportunities for pollen immigration; gene flow is unlikely to contribute substantially to the patterns observed among populations or years.

## Conclusion

We found evidence that all four *N. menziesii* populations demonstrated adaptive capacity and realized adaptive responses to selection. Environmental change between years lowered population 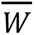 at three of the four sites examined here, and likely influenced the extent of adaptation realized in the progeny generation of our two-generation field experiment. However, despite these strong effects of interannual environmental change, we detected adaptive evolution and tolerance to environmental change in all populations.

All populations experienced substantially different environmental conditions in the second year of the experiment (wetter and cooler than the first), but they differed in how mean fitness changed between years. In some populations, mean fitness increased under the wetter conditions, whereas in others it declined, indicating that the ecological effects of environmental variation differed among populations. At the same time, the realized evolutionary response also depended on environmental context, with genetically based increases in mean fitness occurring only under certain population-by-year combinations. Under a changing climate, populations that harbor substantial V_A_(W) may exhibit genetically based increases in mean fitness that partially offset environmentally induced reductions in fitness. However, further research is needed to determine whether such buffering can sustain population persistence under increasingly intense environmental fluctuations or if directional environmental change may eventually outpace the tolerance limits of populations (Chevin et al., 2010; Milles et al., 2023). Our findings indicate that additive genetic variance for fitness represents the capacity for adaptive evolution, but that the rate at which adaptation is realized is shaped by changing environmental conditions. Determining how evolutionary potential interacts with temporal environmental variation will be essential for predicting the responses of natural populations to ongoing environmental change.

## Author Contributions

S.J.M. conceived of and obtained funding for this work. D. E. G., L. K., H.E.P., and S.J.M. designed the experiment, supervised and conducted the breeding of source seeds, established field sites, trained participating UCSB undergraduate students, collected greenhouse and field data, and interpreted the analyses. H.E.P. performed data analysis and interpretation with the help of A.R.P.. H.E.P. led the preparation of the manuscript, with S.J.M. and A.R.P. revising and editing the text. All authors reviewed and approved the final version of the manuscript.

## Data Availability

Data supporting the findings of this study are publicly available in Dryad￼ at https://doi.org/10.5061/dryad.pvmcvdp1p.

## Code Availability

Code used for analyses and figure generation is publicly available in Zenodo at https://doi.org/10.5281/zenodo.20386658.

## Acknowledgements

We would like to thank Amber Eule-Nashoba for assisting with the initial experimental design (cf. to previous work by Ruth G. Shaw) and for collecting seeds from field populations. We are also grateful to Ivana Gomez, who participated in field work and in the breeding of generation 1, and our 2023 field technicians, Lynn Jung, Riki Radliff, and Ally Wetzel. We also thank Cameron Hannah-Bick for her invaluable assistance with rearing *N. menziesii* seeds in the greenhouse. We greatly appreciate and acknowledge the dozens of undergraduate researchers at UCSB who contributed to greenhouse work, sample processing, and data entry and proofreading. We are particularly thankful to NSF for supporting the following students on Research Experiences for Undergraduate supplements: Michelle Colvin, Bergen Foshay, Cayenne Gularte, Lynn Jung, Jonathan Lam, Grayson Prater, and Rishima Tewari, and for supporting Karen Presburger and Greg Schiller on a Research Experiences for Teachers award. We also extend our sincere gratitude to Ruth Shaw for her invaluable guidance and support throughout the course of this project.

