## Supplementary material for "Estimates of the rate of adaptive evolution in four wild populations of *Nemophila menziesii*"

**Supplemental Materials for "*Inter-annual Change in Mean Fitness in Four Pedigreed Populations of the California Wildflower, Nemophila menziesii: phenotypic plasticity vs. genetic change*"**

**<u>Table of Contents</u>**


**Supplementary Protocols** **1**

S1. Seed Cultivation and Greenhouse Conditions 1

Table S1. Sowing dates and corresponding greenhouse environmental conditions for each experimental cohort. 3

Table S2. Fruit set in the greenhouse in the absence of pollinators. 3

S2. Anther Removal 4

S4. Seed Sterilization 7

S5. Seed Planting 10

S6. Fruit Collection 11

S7. Fruit & Seed Counting 12

Figure S1. Labels for fruit collection envelopes. 14

Figure S2. Identifying filled and unfilled seeds. 15

Figure S3. Glassine envelope labels. 15

S8. Seed Inspection 16

S9. Seed Weighing 17

**Supplementary Figures:** **18**

Figure S4. Climatic variation across California at AC, BB, BO, and HR sites. 18

Figure S5. Mean air temperature (°Celsius) at each of the four field sites. 19

Figure S6. Cumulative rainfall (mm) at each of the four field sites. 20

Figure S7. Experimental design at field sites 21

Figure S8. Aster graphical model used to estimate W for each generation at each site. 22

Figure S9. Bivariate Spearman rank correlation for estimated sire effects on W of parental family groups sown in 2022 and 2023 for the field sites AC, BB, BO, and HR. 22

Figure S10. Probability densities of sire effects on fitness for the estimated mean lifetime fitness of parental sibships between years. 23

**Supplementary Tables:** **23**

Table S2. Summary of sibships, transects, and seeds planted in generation 1. 23

Table S3. Population structure and seed planting overview for Fall 2022. 24

Table S4. Parameter estimates from generation 1 and 2 aster fitness models. 24

Table S5. Likelihood ratio tests evaluating genotype-by-year (G × Y) interactions for fitness within each population. 25

Table S6. Seed size and survival to flowering. 25

Table S7. Mean seed size by generation. 26

### Supplementary Protocols

#### S1. Seed Cultivation and Greenhouse Conditions

In August 2018, seeds were planted individually into moist coconut coir (COIR) plugs and stored in a dark, cold room maintained at 9°C to promote stratification and germination. Due to limited greenhouse space for the adult plants, a total of seven successive cohorts were sown, cultivated, and hand-pollinated. In each cohort, we raised and pollinated 7-10 mating groups representing each population. As individuals flowered, they were randomly assigned to mating groups for logistical purposes. Mating groups were used to organize pollinations and corresponded to half-sibling groups.

Each cohort was comprised of ~350 seeds representing ~43 field-collected maternal families from each of the four populations, with the exception of cohort 7, which required seeds from only Angelo Coast Range Reserve and Hastings Natural History Reservation in order to achieve the planned sample size. Male-sterile plants were always assigned as pollen recipients. After one month, the plugs were removed from cold storage, and any mold-contaminated seeds were discarded. Germinated seedlings were transplanted into 4-inch pots and placed in refrigerated, open-top shelving (maintained at 10-15°C) in a greenhouse. Once well-established, the seedlings were transferred into 6-inch pots to promote continued vegetative growth prior to flowering. From each population, we aimed to maintain two flowering individuals per maternal family to ensure that one individual per family would be available for hand-pollination.

In each cohort, as plants began to flower, they were moved from the chilled room to a warmer greenhouse for breeding. The greenhouse temperature was regulated automatically, with heating activated when temperatures dropped below 18.3°C and cooling engaged when temperatures exceeded 26.6°C. Individuals representing a given population were located on a large rolling table, with each population located in a different section of the greenhouse. Within each cohort and each population, to create a nested breeding design, ~10 plants (all producing healthy pollen) were selected at random to function as males (pollen donors) and the rest as females (pollen recipients). Within each population (in each cohort), each of the pollen donors was assigned three pollen recipients at random. In each population, the pollen donors were arrayed in a single column, with pollen recipients were arrayed in random locations on either side of this column. Each of the individuals used in this nested half-sib design was derived from a different field-collected maternal family.

The greenhouse was not large enough to accommodate all individuals to be pollinated, so we raised and pollinated a subset of the field-collected maternal families from each population in seven successive cohorts from August 2018 to December 2019 (see supplementary material for details). In each cohort, from each population, we cultivated two seeds representing each of approximately 40 field-collected maternal seed families. Upon flowering, these plants were assigned at random to individual mating groups, each composed of one pollen donor and three pollen recipients derived from different field-collected maternal families. To the extent possible, only one seed from each field-collected maternal family was used. However, to ensure that each mating group included co-flowering individuals, in some cases, the two seeds cultivated from a maternal family were required to be used. In such cases, one of the two offspring was used as a pollen donor in one mating group, and the other was used as a pollen recipient in a different mating group. The number of maternal seed families used for both donor and recipient roles (i.e., two differed among populations (AC: 18, BB: 3, BO: 21, HR: 34). In no case was the same maternal family used as a pollen donor more than once, or as a pollen donor and as a pollen recipient in the same mating group. In each population, across all cohorts, ~50 individuals were designated as pollen donors (sires) and 150 as pollen recipients (dams).

Controlled hand pollinations were conducted using fine paintbrushes and jeweler's glasses to ensure accuracy. Flowers on the pollen recipients were emasculated prior to anthesis to prevent self-fertilization, and pollen was applied manually to receptive stigmas from designated pollen donors. During cohort 4, emmasculated plants were embedded throughout the experiment to determine if pollination was occurring due to insect pollinators in the greenhouse. A near-zero rate of fruit set confirmed that unintended pollination did not occur (N = 1,651 flowers, 0.02%). The pedigreed seeds produced from these crosses were subsequently stored under cold conditions (9°C) until planting in 2021.

**Table S1.** Sowing dates and corresponding greenhouse environmental conditions for each experimental cohort.

The table lists the date each cohort was sown.

| Cohort | Date sown |
|---|---|
| 1 | Aug 7th, 2018 |
| 2 | Sept 14th, 2018 |
| 3 | Dec 18th, 2018 |
| 4 | Febr 12th, 2019 |
| 5 | March 19th, 2019 |
| 6 | April 26th, 2019 |
| 7 | July 27th, 2019 |

**Table S2.** Fruit set in the greenhouse in the absence of pollinators.

Emasculated dams were placed throughout the greenhouse among the hand-pollinated pollen recipients during the period of hand-pollination and seed development. These plants were monitored during cohort 4 to determine the rate of fruit set in the greenhouse due to either self-pollination or the presence of unintended pollinators. N = 1651 flowers, of which 32 set fruit. Fruit set was 0.019%.

| Population | Dam ID | # of aborted flowers | # of ripe fruit |
|---|---|---|---|
| AC | 63 | 116 | 0 |
| BB | 84 | 125 | 1 |
| BO | 166 | 162 | 2 |
| AC | 57 | 168 | 0 |

| | | | |
|---|---|---|---|
| BB | 153 | 103 | 0 |
| AC | 58 | 53 | 0 |
| HR | 36 | 190 | 1 |
| BO | 127 | 113 | 4 |
| HR | 88 | 194 | 1 |
| BB | 61 | 61 | 4 |
| HR | 182 | 181 | 0 |
| BO | 3 | 55 | 1 |
| BB | 71 | 98 | 18 |

**S2. Anther Removal**

1. Wash hands with soap and dry them with a clean paper towel.
2. Assemble the supplies and datasheet
3. Sit at a comfortable height in front of the plant to be emasculated.
4. Double-check the Plant ID of the plant by inspecting the plastic label in the pot.
5. Identify the correct row (with the same Plant ID) in the data sheet on which to record the number of flowers emasculated.
6. Make sure to write your initials on the row associated with each plant, emasculate
7. Clean forceps (both the inner and outer surfaces) with a paper towel that has been sprayed with ethanol (EtOH).
8. *Perform on one bud or flower at a time*: Remove the anthers.
9. As you remove the anthers from a flower, count them to make sure that no anthers remain inside the flower. The vast majority of flowers produce 5 anthers, but occasionally a flower will produce 6 anthers, so make sure all anthers have been remmoved.
10. After anther removal from a flower, place a label (with the correct color coding the date) around the pedicel of the flower. The "pedicel" is the stalk that attaches a flower to the stem. Do not place the label on the stem. The label must be on the pedicel.
11. Remove the anthers from up to 10 closed buds and open flowers on the focal plant.
12. As each bud or flower on pollen recipient is emasculated, keep track of the total number emasculated.
13. Record the number of buds and flowers that are emasculated on each pollen recipient in the proper column in the Emasculation data sheet.

Tips:

- If the anthers of a flower on a Dam are already releasing pollen (if any pollen has dropped onto the petals), then the flower should be removed. Using the fingers of one hand, fold the petals of the flower towards the center so that no pollen is deposited on your fingers. With the other hand, hold the pedicel while pulling off the flower (don't pull off the flower with only one hand, as this can cause the stem to break!). Place the discarded flower in a small container that you bring with your from plant to plant.
- When emasculating closed buds, gently twist the corolla (the petals) between two fingers in order to force the bud to open slightly; be careful not to detach the bud from its pedicel.

- Once the bud has begun to open, insert the closed tips of the forceps between two petals in order to force them to open up, exposing the anthers. Only when the bud has been forced to open a bit should the forceps' tips be enabled to separate. At that time, the forceps may be used to grip a stalk of an anther (the filament) in order to remove it from the flower.
- Be very careful not to grab a stigma or the style because if you pull on these organs, this will damage the flower such that it cannot be pollinated..
- Always use a pencil to record data on the data sheet.
- Write neatly. 7's must be crossed. 1's must be very legible. 9's must have a closed loop. 4's must NOT have a closed loop because it makes them look like a 9.

**S3. Pollination**

1. Wash hands with soap and dry them with a clean paper towel.
2. Assemble the supplies and datasheet
3. Use the data sheet (and clipboard) for the population to be emasculated.
4. Write today's date on the data sheet
5. Locate and retrieve the index card that contains the locations and the Plant IDs of the Sire and the Dams in the sibship to be pollinated.
6. Note the labels that identify each tabletop in the greeenhouse.
7. Each table is divided longitudinally into two halves. The "A" side is on the left of each table, and the "B" side is on the right of each table.
8. The length of each side of the table is divided into numbered positions, with position 1 closest to the greenhouse entrance. The positions increase sequentially along the edge of the table, progressing from the entrance (or front) of the greenhouse to the back of the greenhouse.
9. When preparing to pollinate a given mating group, first visit the Sire and record (in the column adjacent to the Sire's ID) the total number of flowers that are open and producing enough pollen that you could use them for pollinations. If there are >10 pollen-producing flowers, simply write ">10".
10. Then, visit each of the 3 Dams in the mating group (these are the plants that you will pollinate) and record the number of flowers that are ready to be pollinated on each Dam. A flower is ready to be pollinated when its petals are fully open and the stigma branches have diverged and are arching outwards; in addition, the stigma tips must be dark blue and visibly swollen. These stigmas are said to be "receptive".
11. Add the total number of flowers that are ready to be pollinated in the mating group.
12. Return to the Sire of the mating group that you are pollinating. For every 3-4 flowers that need to be pollinated in this mating group, collect 1 pollen-producing flower from the Sire of that mating group. Place the pollen-producing flowers in a weigh boat (one may return to the Sire to collect more pollen-producing flowers if needed and available).
13. Carrying the weigh boat with the pollen-producing flowers and the clipboard, return to each of the 3 Dams to pollinate them. If there are not enough pollen-producing flowers to pollinate all of the receptive stigmas, then divide up the pollinations among the Dams. For example, if only 2 pollen-producing flowers are available from a mating group's Sire, but there are a total of 5, 3, and 7 flowers (15 total) that are ready to be pollinated on the 3 Dams, then pollinate 2, 2, and 3 flowers (7 total) on the Dams with the available pollen.
14. Double-check the Plant ID of each Dam (prior to pollinating it) to make sure that it is supposed to be pollinated by the Sire whose pollen you collected
15. Seat yourself at a comfortable height in front of the Dam to be pollinated.
16. Holding the pedicel of a pollen-producing flower, bring it "face to face" with a flower bearing receptive stigmas on the Dam to be pollinated. Brush an anther that is releasing its pollen across the surface of the stigma. The pollen that is transferred to the stigma should be visible as the stigma surface becomes lightly covered with a layer of cream-colored pollen. A single anther should produce enough pollen to cover the surface of both stigma tips in a given flower.
17. After a flower has been pollinated, gently place the pollen-producing flower back into the weigh boat and then write the letter "P" on the label of the pollinated flower.

18. *Keep careful track of the total number of flowers that you have pollinated on a given Dam and record this number in the data sheet*. Some students like to keep an index card with them to make a mark for every flower that they pollinate on a Dam and then add up the marks in order to record the total number of flowers pollinated.
19. Write your initials in the correct column and row of each Dam that you have pollinated.

Tips:

- Use the magnifying visors to verify that the stigmas in a flower are ready to be pollinated.
- Use the magnifying visors to convince yourself that you have successfully applied pollen to a receptive stigma.
- Always use a black "Artists" pen to write the "P" on the label of every pollinated flower.
- Always use a pencil to record data on the data sheet.
- Write neatly. 7's must be crossed! 1's must be very legible. 9's must have a closed loop. 4's must NOT have a closed loop because it makes them look like a 9.
- If the anthers of a flower on a Dam are already releasing pollen (if any pollen has dropped onto the petals), then the flower should be removed. Using the fingers of one hand, fold the petals of the flower towards the center so that you don't get any pollen on your fingers. With the other hand, hold the pedicel while you pull off the flower (don't just pull off the flower with one hand). Place the discarded flower in a transparent plastic basket that you bring with your from plant to plant.

**S4. Seed Sterilization**

Preparation

- Clean off workspace on bench with 70% Ethanol or Clorox wipe
- Clean 2 stainless steel spatulas per person with 70% Ethanol; keep them off the surface by placing them horizontally (when not in use) on clean weigh boats. Wash your hands!

Amount of solution needed depends on the number of seed samples (envelopes) to be cleaned:

| Number of samples to be sterilized | 10% bleach<br>1% Tween<br>89% purified H2O<br>40-mls poured over seeds through filter paper | Water rinse<br>(40 mL x 5 times)<br>Purified H2O, boiled and cooled to room temperature in ice-slurry bath | Time required (approx.) for prep & treatment<br>1 bleach rinse + 5 water rinses, then transfer seeds to petri dish lined with filter paper |
|---|---|---|---|
| 13 | 520 ml | 2600 (13 x 40 mL x 5) | 90 mins |
| 25 | 1000 ml | 5000 | 2 hr 55 min |
| 50 | 2000 ml | 10000 | 5 hr 45 min |

<u>Supplies:</u>

Water for rinsing:

Two 2800-ml Erlenmeyer flasks for cooling water

Kettles with DI water to fill beakers ¾ of the way full

For bleach solution (recipe below):

Clorox bleach

Tween

Purified water

Other things:

Scissors for opening seed envelopes

Petri dishes (10 cm diam; tops only) – one for each sample

Number 1 filter paper (15cm diam) for glass cones

25 Erlenmeyer Flasks (1000-mL & 500-mL)

25 glass cones (long- and short-stemmed)

6” deep bin, filled with ice and water slurry to cool purified water after boiling

Seed envelopes containing glassine envelopes with viable seeds

2 x 50 mL graduated cylinders: one for only bleach solution & for only boiled purified H20

Preparatory Tasks

<u>Mix Bleach Solution:</u>
Prepare (rinse with purified water) 2 one-liter bottles with tight screw cap each to contain:
100 mL bleach
890 mL DI water (no need to boil)
10 mL Tween

Important: Tween will be viscous, so mix using stir-plate (with magnetic stir rod). Add the Tween last and let it drip for 1-2 min. After mixing, cap tightly and shake gently to homogenize.

Prepare Purified Water

Boil DI water in clean electric tea kettles for one minute (or microwave in 1 L beakers).

Rinse two 2800 mL E. flasks with boiled water, then fill them with the boiled water.
Place flasks in ice water slurry. Agitate every few minutes to facilitate cooling. Divide the boiled water among smaller beakers to cool faster. Tip: Prepare boiled & cooled purified water ahead of time.

<u>Sort & Store Dried Seeds:</u>
Assemble the empty seed envelopes of the seeds that have been drying for at least 5 days.

Place these in a line along the edge of the large bench, farthest from the lab entrance.

Carefully place petri dishes containing dried seeds next to their associated envelopes on the bench (double check both the population and maternal ID).

Transfer dried seeds into newly labeled glassine envelopes (with population, maternal ID, and ‘G2’). Tape the bottom edge of the new glassine envelope before folding and storing it in the yellow seed envelope.

Sort completed envelopes into clean ziplock bags, then into boxes labeled by population. Fill the bags with ample silica gel and seal them. Move the box to the cold room when the population is complete.

Bleaching Prep:
If two people are working on this task, then one person should start at one end of the line of beakers , and the other should begin in the middle, with both people working in the same direction.

1. Assemble seed envelopes to be treated in this session (plan to treat 25 at a time).
2. Prepare glass funnels, filters, and beakers. Beakers (with their funnels) should be arranged in grid cells ~1.5 feet apart along the edge of the large black bench.
3. Fold #1 filter papers (15 cm diam.) into quarters and place one in each glass funnel cone
4. Prepare 25 petri dishes with 9 cm diameter filters (one lines the bottom of each dish).
5. Place one seed envelope and its assigned petri dish in front of each filtration set-up.
6. In pencil, along the edge of each petri dish filter paper, write the Population & Maternal ID (e.g., BO_099) and date of bleaching (mm/dd/yr) (e.g., 10/11/22) for the envelope.
7. Carefully place the seeds from the glassine envelope into its assigned glass filter cone lined with a folded filter: Keep filter paper fully open to avoid seeds slipping between folds.
8. Double-check the glassine & envelope for any remaining seeds. Make sure the petri dish paper is labeled correctly. Move petri dish and envelope next to the beaker, out of the "splash zone".

Bleaching:

9. Pour 40 mL of bleach solution into each seed-containing filter cone. Pour slowly so that the solution doesn't overflow and seeds aren't lost. If any seeds are floating, brush them gently to the surface of the filter cone to dislodge air bubbles. This should cause them to sink.
10. Place bleach solution bottle with tightly closed caps on work bench.
11. Seeds should remain in bleach for 5 min before rinsing (keep the time from the first pour).

Rinsing:

12. After the boiled, purified water has cooled, pour 40 mL into each glass cone using the same slow pour used for bleach. If two people are working, both can decant water into two narrow-mouthed 500-mL beakers, and each person can rinse seeds independently with 40-mL aliquots of water.
13. After each sample has received 40 mL of cool, boiled water, repeat the process until each sample has been rinsed five (5) times with 40 mL of purified water each.
14. Carefully remove the filter paper holding the seeds from the glass cone and unfold it. Using the flat side of a clean metal spatula, transfer each group of seeds to the single petri dish with filter paper labeled with the correct population, maternal ID, and date.
15. Gently spread out the seeds into a single dispersed layer on the filter paper. Use a spatula or tap dish on the table. No seeds should be touching.

16. Arrange petri dishes in a green tray with 4-5+ cm of space between dishes. Cover each petri dish. Place trays in a fume hood (in large or small lab) for 5-7 days to dry, ensuring no tray is completely covered. As seeds and the filter paper dry, they will be more likely to jump around.
17. Organize the empty seed envelopes from this seed-sterilization session into a separate small container. With tape or a sticky note, label the date on which seeds from these envelopes were bleached. Keep these envelopes separate from any other seed-containing envelopes until the seeds are transferred back into them.

Down Time Tasks

- Boil more purified water and allow to cool to increase rinse water stocks.
- Prepare more bleaching solution.
- Clean & sterilize petri dishes for the next bleaching session
- Empty and clean glassware at filtration stations
- Sort and store dried and bleached seeds

**S5. Seed Planting**

To ensure accurate identification and monitoring of planted seeds, we used a combination of peat pots and labeled skewers. Peat pots were inserted into small excavated holes prior to sowing, which minimized the likelihood that seeds from the existing seed bank would germinate within the same location and be misidentified as experimental individuals. The peat pots also reduced seed displacement caused by rainfall between planting and subsequent censuses. Labeled skewers were used to mark the position of each planting location, facilitating relocation in the field. While peat pots were not expected to increase germination success relative to direct sowing, their use, in combination with skewers, improved the accuracy of tracking individual seeds and seedlings over time.

1. *Preparation of Trays with Peat and Coir Plugs:* Aero 210 trays produced by QuickPlug NA Inc. (quickplug.com) containing peat and coconut fiber (coir) plugs were labeled using duct tape to indicate population and treatment information. Each tray contained 210 plugs, 1.6 centimeters diameter and 3.3 centimeters tall. A sanitized tool was used to create a single divot in each peat plug to receive seeds.
2. *Seed Placement and Cold Storage:* Seeds were removed from glassine envelopes and placed individually into the prepared peat plug divots. Trays were then transferred to a cold room maintained at 9 °C for stratification prior to field planting.
3. *Pre-Planting Storage and Transport:* Seeds were stored in the cold room for 24–48 hours before field deployment. During transport to the field site, trays were kept covered and handled carefully to minimize disturbance and maintain moisture.
4. *Field Site Preparation:* At the field site, 1 m segments were cleared of existing vegetation. Within each segment, 8–10 planting holes were created using an apple corer to ensure consistent depth and spacing.

5. *Transplanting and Identification:* Peat plugs were carefully removed from trays and placed into the prepared planting holes. Each planted plug was marked with labeled skewers and flags to identify individual plants and distinguish them from their source population.

**S6. Fruit Collection**

*Nemophila menziesii* Fruit Collection in the Field
Spring 2023 Protocol

1. After confidently identifying the fitness plant, locate all branches. Apical and lateral branches may be long and windy (as at Bodega Bay) or short and unbranched. Be careful not to damage the plant when finding branches!

2. Gently lift each branch to check the underside for fruits. *N. menziesii* pedicels curve downwards as fruits develop, so some fruits may even be buried under 1-2 cm of soil.

3. Identify and collect closed near-mature and mature fruits. Near-mature fruits will appear mostly green, sometimes with blotches of purple or other dark coloration. Fruits should be swollen to the point that the calyx (sepals) have started to be pushed out. Closed mature fruits may appear non-green (purple to brown) in color and be noticeably more swollen/larger than near-mature fruits. The calyx of these fruits will turn from green to yellow or brown, becoming more brittle and dry. In both cases, pedicels will be more curved and often more rigid. Also, collect any closed fruit when, after *gently* squeezing it between your fingers, it pops open!

    a. You should pinch fruits off below the receptacle, such that a portion of the pedicel remains attached to the fruit upon collection.

4. Collect all open fruits. Any fruit with noticeable dehiscence should be classified as an open fruit! Unless it is completely closed before we collect it, it's open. These should always be collected in coin envelopes separate from closed fruits.

    a. You should collect all closed fruits on a plant before moving on to collect open fruits.

5. Label fruit collection envelopes in pencil with the date, population & recipient ID, Point/Transect, and Plant number. Population + Recipient ID can be written with the population code, followed by a '–' or 'x' for generation 1 or generation 2, respectively. Write the Recipient number with three digits (including leading zeros)!

    For example, closed fruit from a parental fitness plant:

    HR – 067 4/04/2023
    Tr. 150
    plant 5

F-10

Closed

---

a. Write as integer counts (not tallies!!) the number of fruit collected below this label information. E.g.: F-24

b. Write “closed” or “open” near the bottom of the envelope to indicate the type of fruits collected.

6. Seal the envelope once all fruits of that type have been collected from a plant. Double check that the envelope has been labeled correctly.

7. Place completed fruit envelopes into a ziplock bag containing ~1/3 cup of new/re-desiccated silica beads. Label this bag with the population & collection date.

**S7. Fruit & Seed Counting**

Preparation:
- Wash your hands thoroughly with soap and water!
- Prepare your work station by cleaning the bench surface. Make sure you have the following supplies:
  - 70% ethanol
  - Kimwipes
  - ≥ 4 large weigh boats (clean)
  - Forceps of different sizes (also clean)
  - Glassine envelopes
  - Scotch tape
  - Pitt pen

  You should be working on a surface of 2-3 pieces of clean printer paper (or other light-colored paper) taped to the bench surface. This will allow any escaped seeds to stand out due to the higher contrast.

Note: Work with one population at a time. Work through collection (coin) envelopes following the sequence of Maternal IDs within each transect/point. Envelopes should be organized by the following scheme: Closed vs Open fruit → Transect/Point → Segment Number→ Left or Right → Collection Date
All fruits collected will be from fitness plants, so there should *almost never* be multiple plant IDs for a given Maternal ID x Transect x Gen combination.

<u>Fruit Counts</u>

1. Pull out the next envelope to have its fruit counted from the collection box, following the order described above. Note whether the envelope you pull contains “CLOSED” or

"OPEN" fruits. Also note the date the collection was made. Open the envelope carefully, using scissors if necessary. Work only with one envelope at a time, and keep track of where you place the envelope!

2. Carefully empty the contents of the envelope into a clean weigh boat. Make sure no seeds are stuck in the bottom crevice or corners of the envelope. Count the number of whole fruits collected. Count them again, and double-check that the count you obtain matches the one written on the envelope.
    a. If your count matches the written count, write your initials on the envelope directly next to the fruit count in red marker (e.g., "DEG").
    b. If your count does not match the envelope, cross out the envelope count and write the true count in digits (not tallies!) directly next to it in red marker. Add your initials next to this true count*.

    *Note: A whole-fruit is determined by the presence of the receptacle and at least half of the ovary (fruit) wall). Fruits may have broken up into pieces following collection (especially OPEN fruits, which will be trickier to count in general), so get a second opinion if you're unsure whether the fruit you have is whole or unsure about the total count.
3. Record the true total fruit count in the corresponding row and column of the data sheet (CLOSED or OPEN), double-checking the envelope's label information against the collection date, transect, maternal ID, generation, segment, and plant number in the data sheet. Note: dates that are one day off from the recorded collection date were collected on the same trip and should be entered under the same column. Be sure to also double-check the fruit type (closed vs open) and the count you entered!
4. Proceed to the steps below for counting the number of viable seeds only if the fruits in the envelope were CLOSED when collected. If working with OPEN fruits, return the entire contents back to the envelope it came from. Tape this envelope closed and return it to its proper place in the fruit collection box.

<u>Seed Counts</u>

Only count the number of viable seeds from fruits that were CLOSED when collected. These counts will give us a more reliable estimate of the average number of seeds produced per fruit by the plant. Check that you have followed all the preparatory steps as described above.

1. Using an additional one or two clean weigh boats, separate the seeds from their fruits and place them in their own weigh boat. Remove any other fruit tissue and debris from this weigh boat so that it contains only the seeds. Double-check each individual fruit for seeds that may be stuck inside the fruit wall.
2. Remove the light-colored placentas of all the seeds using forceps and retain only the "de-placentaed" seeds in the weigh boat.
    a. The placentas, empty fruits, and non-seed debris can be discarded or moved to a "discard" weigh boat (to be emptied into a trashcan when full).
3. Carefully move the weigh boat of seeds to a dissecting microscope. Separate the "filled" seeds (those appearing fully "inflated") from the unfilled seeds using forceps or a spatula.
    a. Visually inspect each seed to determine to the best of your ability whether that seed is "filled" (likely viable) or "unfilled" (likely inviable). Filled seeds are rounder and fuller without many flat or jagged edges. Filled seeds are usually (but not always!) a bit larger and darker than unfilled seeds (*see images below*).

    b. Separating seeds into three different groups may help: unfilled, filled, and unsure.
    c. Microscope alternative: Use the head visors (x 3.5 mag) at your station. If you're using the head visors, please make sure that there is enough light to give you a good view of the surface of the seeds.
4. Have a second person check your sorting, and discuss any seeds you're unsure about before deciding on a final filled seeds count. If necessary, have a third person check particularly ambiguous seeds.
5. In red ink, write the number of filled seeds ("# filled") you counted below the fruit count on the collection envelope as a fraction of the total number of seeds, followed by your initials. Record this count in the appropriate cells of the data sheet, double-checking the envelope's labeled information against the transect, maternal ID, generation, plant number, and collection date in the data sheet. Also, record the total number of seeds (filled + unfilled) in the data sheet to the right of the number of filled seeds.
    a. Double-check that all information is entered correctly!!
6. Return all of the unfilled seeds directly to the yellow collection envelope they came from. Transfer the filled seeds to a new glassine envelope labeled with all of the info present on the yellow envelope (except # fruits and closed/open), plus the number of filled seeds/ total number of seeds, writing in a black Pitt pen on the front. Tape the bottom of the glassine envelope with scotch tape to make sure seeds cannot escape. Close the glassine envelope, then fold it over itself several times before placing it in the yellow collection envelope. Tape the coin envelope shut with a piece of scotch tape. Return the envelope to the box it came from.

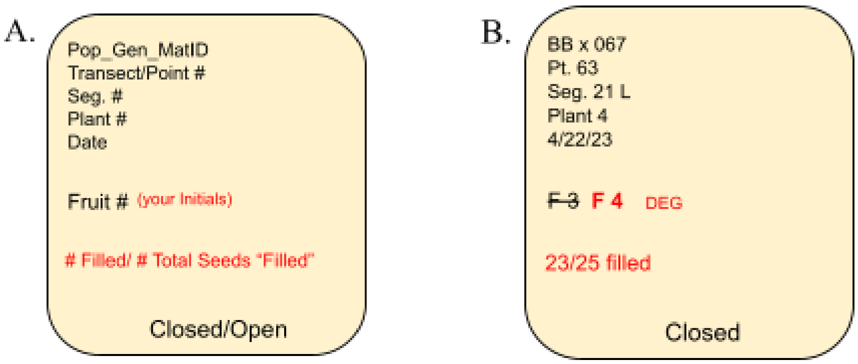


**Figure S1.** Labels for fruit collection envelopes.

A) A template for how fruit collection envelopes should be labeled. B) An example of such an envelope for closed fruits collected at Bodega Bay, with the original fruit count corrected.

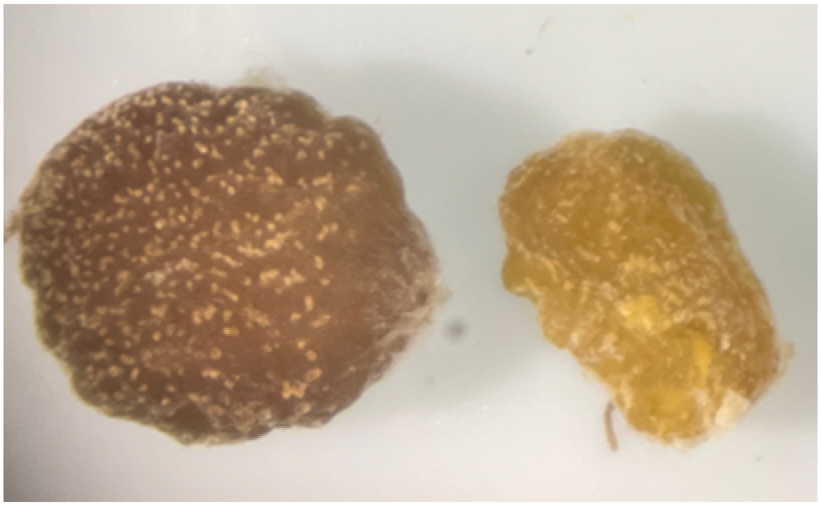

**Figure S2.** Identifying filled and unfilled seeds.

A filled seed (left) next to an unfilled seed (right). Note the differences in shape, size and color. Also know that this example is more idealistic than what we often see: there is lots of variation within each of these categories.

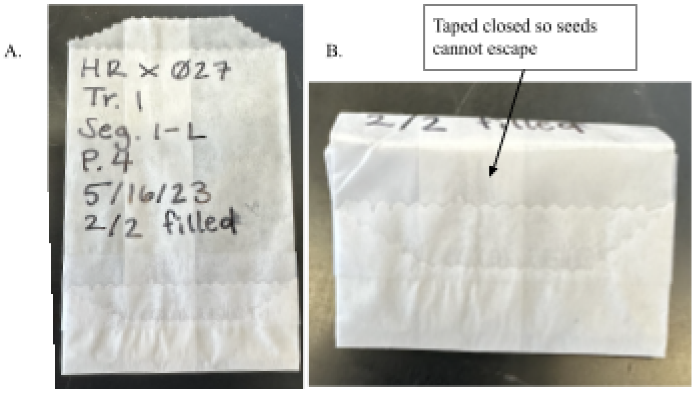


**Figure S3.** Glassine envelope labels.

An example of how the glassine envelope should be labeled with the maternal ID, generation, transect/point, segment, plant ID, date, and filled number/total number of seeds. Glassine envelopes should be taped to ensure that seeds cannot get lost.

<u>Seed Weights</u>

We will weigh batches of filled (viable) seeds to obtain their total weight in milligrams. Check that you have followed each of the preparatory steps described at the beginning of this document.

Additional Preparation:

- Clean the bench surface in front of the OHAUS microbalance scale. You may tape white paper onto the surface to aid in spotting escaped seeds
- Obtain two small clean weigh boats and clean forceps, spatula

- Turn on the OHAUS microbalance scale and make sure it is balanced and properly zeroed

1. Extract the glassine envelope with filled seeds from the next yellow coin envelope. Pour these filled seeds into a clean weigh boat and COUNT them to make sure that they match the number of filled seeds written on the envelope and recorded in the datasheet. Also, check that all the other identifying information on the envelope (including date) matches up with the row and column in the data sheet.
    a. If the seed count you get does not match the previously recorded count, consult a graduate student or REU student before making any changes to the data
2. Tare a clean small hexagonal weigh boat on the scale.
3. Carefully transfer 30 filled seeds per envelope (or as many as available if fewer than 30) to the tared weigh boat. If there are more than 30 seeds, choose 30 randomly to weigh. Make sure all the placentas are removed from the seeds. Place this weigh boat on the scale. Record the number of seeds being weighed in the datasheet in the appropriate row. If there are zero-filled seeds, input "0" for the seed count and "N/A" for the seed weight.
4. Wait 60 seconds for the scale to settle on a weight. Then, record the total mass of the seeds weighed in the datasheet. Double-check that you're entering data in the proper row and that there are no typos in the number of seeds weighed or the total weight.
    a. If the scale is not settling on a weight or it is saying "busy", consult a graduate student or REU student to help clean and recalibrate the scale.
    b. If a seed is lost, note on the envelope that a seed was lost. Make sure the "# of seeds weighed" column reflects the true number of seeds weighed. If more than 30 seeds are available, you may retrieve an additional seed to weigh a total of 30 seeds.
5. Carefully return all filled seeds to the glassine envelope. Fold the glassine over itself several times before placing it back into the yellow coin envelope and taping it shut. Move this coin envelope to a "complete" pile to be returned to the collection box, and move on to the next batch of seeds.
6. Re-tare and clean the weigh boat every few envelopes to prevent contamination.

### S8. Seed Inspection

1. Wash hands
2. Remove the plastic covering on the microscope. Keep close at hand to return later
3. Get 2 weigh boats. One should be labelled "unweighed" and can be found at the microscale station. The other can be any unlabeled and clean weigh boat.
4. Bring the seed envelope with the red number to the seed inspection station. The red number indicates recounted fruits.
5. Clean weigh boats with ethanol. There is a spray bottle with 70% EtOH on the big, black lab table (where the boxes of seeds are), and paper towels next to the sink, which can be used to clean any debris from a weigh boat.
6. Add seeds to the unlabelled seed boat
7. Get tweezers and spatula, and clean with ethanol. Use tweezers to pick up seeds and spatula to move around seeds
8. Focus seeds under microscope. Use lower magnification and coarse tuning to get clear image

9. Look for seeds that are round and don't have indentations. Choose 5 of these seeds and add to weighed seed boat
10. Remove any white connective tissue from your chosen seeds
11. Crush any dried fruits and shake out any seeds if you cannot find enough good seeds
12. Give seed packet, and boat labelled "unweighed" to seed weigher

**S9. Seed Weighing**

Using the Cahn Microbalance:
Before you start a weighing session, you must tare the balance with nothing on the round weighing pan. This consists of the following steps.

1) Wash your hands
2) Assemble clean forceps, plastic hexagonal weigh boat, data sheet, pencil, and weighing paper folded into a weigh boat or the premade aluminum square weigh boat
3) Close the glass window if it isn't already closed.
4) Lower the stabilizing brake (press the brake button to lower the column) to allow the weighing pan to swing freely. WAIT until the pan stabilizes.
5) Press TARE. Wait until the display stabilizes at 0.000
6) Raise the BRAKE (pressing the BRAKE button once more)

You should not have to tare the empty weighing pan again during this session.
To weigh a group of seeds:
Make a stable "weighing boat" out of weighing paper, or use the tiny square aluminum folded tray that's next to or inside the window.
1) Tare the weighing pan *with the weighing boat on it by following these steps*
a) Be sure the brake is raised, supporting the round weighing pan
b) Put the small empty weighing boat on the round weighing pan. Do not under any circumstances touch the wires that hold the weighing pan with either the forceps, the weigh boat, or your fingers.
c) Close window
d) Lower the brake (so that the round weighing pan swings freely
e) Press tare
f) Wait until the display stabilizes at 0.00
g) Put the brake back up (raise it)
h) Open the window and remove the weighing boat
i) Close window
2) Place 10 seeds in the weighing boat.
3) Place the weighing boat on the round weighing pan. Do not touch the wires. Keep your hands stabilized.
4) Close the window
5) Lower the brake so that the round weighing pan (with the aluminum tray) swings freely.
6) Wait until the display read-out stabilizes. Record the taxon, Tag ID, and weight of the seeds in the data sheet. Include two decimal places to the right of the decimal point. The weights are in micrograms.

7) Raise the brake before removing the weighing pan.
8) Open door, remove weighing pan and replace the seeds in their tube.
9) Place the weigh boat with the next group of seeds on the round pan to weigh it. You do not have to tare it between samples.

**Supplementary Figures:**

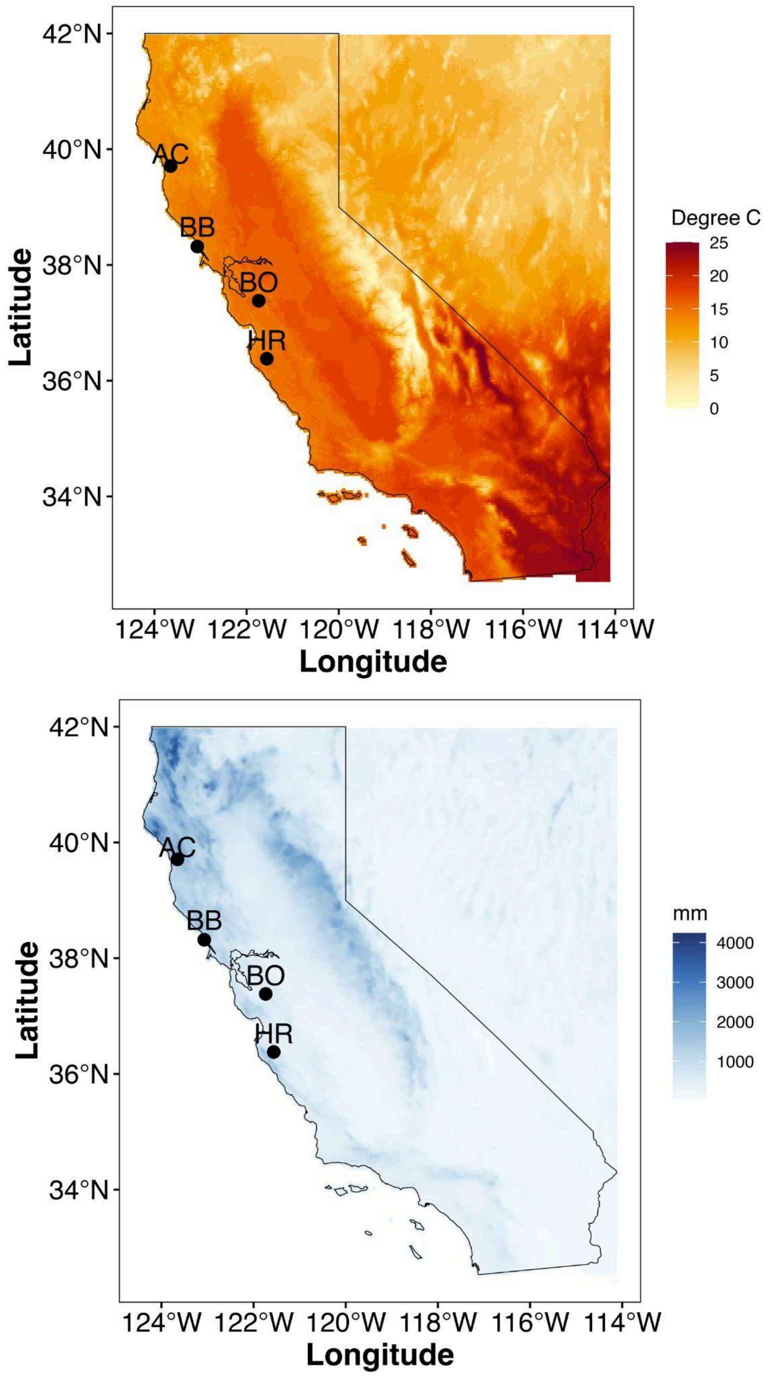


**Figure S4.** Climatic variation across California at AC, BB, BO, and HR sites.

Mean annual temperature (°C) (above), and mean annual precipitation (mm) (below) across California, based on 30-year PRISM climate normals. The locations of the four study populations of *N. menziesii* are marked. These sites span a gradient of temperature and precipitation, with AC and HR representing the northernmost and southernmost extremes, respectively. Temperature increases and precipitation decreases along this latitudinal gradient.

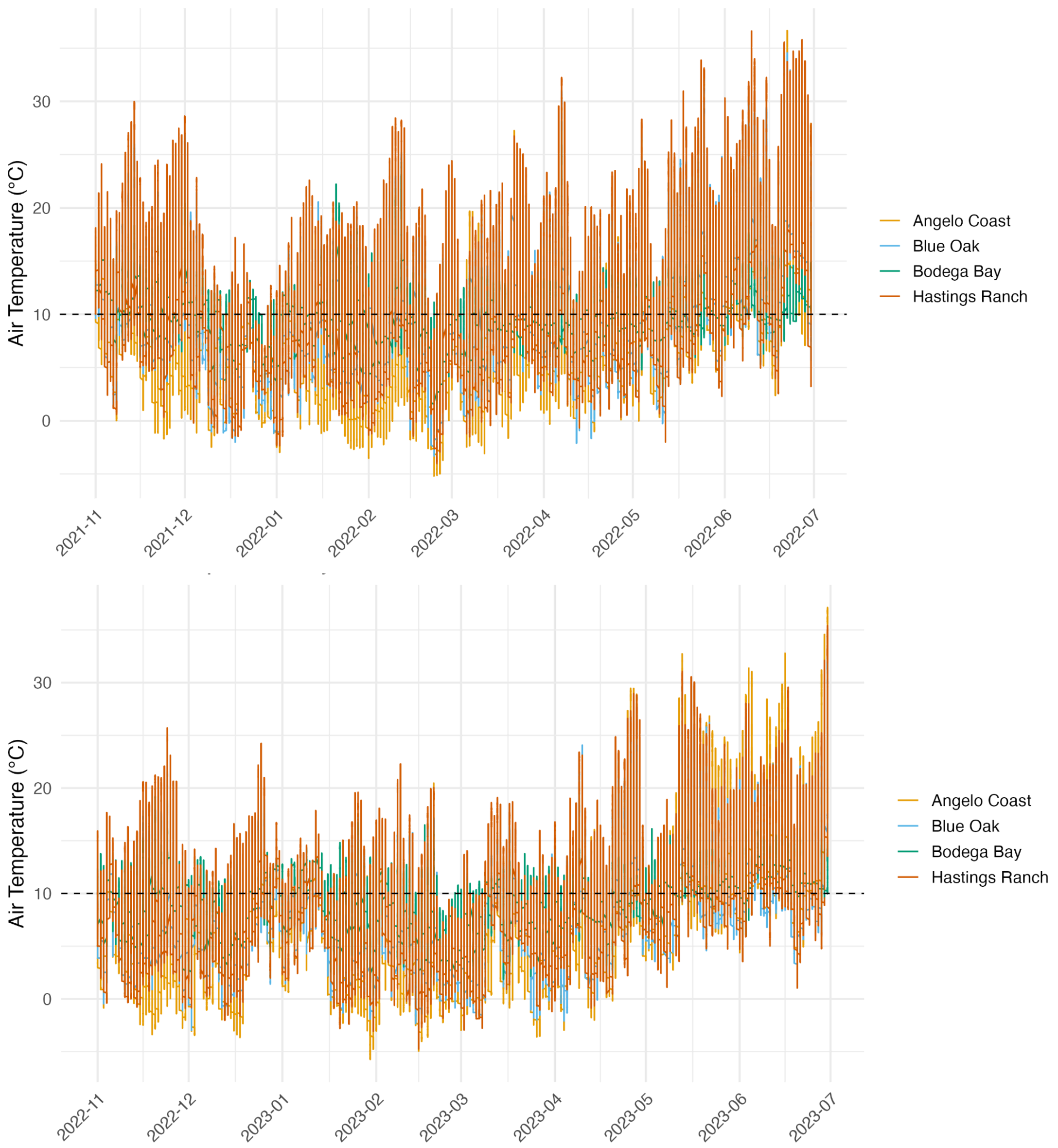


**Figure S5.** Mean air temperature (°Celsius) at each of the four field sites.

Data is collected from mid-December to June for 2021-2022 (above) and 2022-2023 (below) from permanent sensors at each of the UC Reserves. The red line represents 10°C on each graph to aid in comparison. Note that the mean temperature is higher in 2021-2022 compared to 2022-2023.

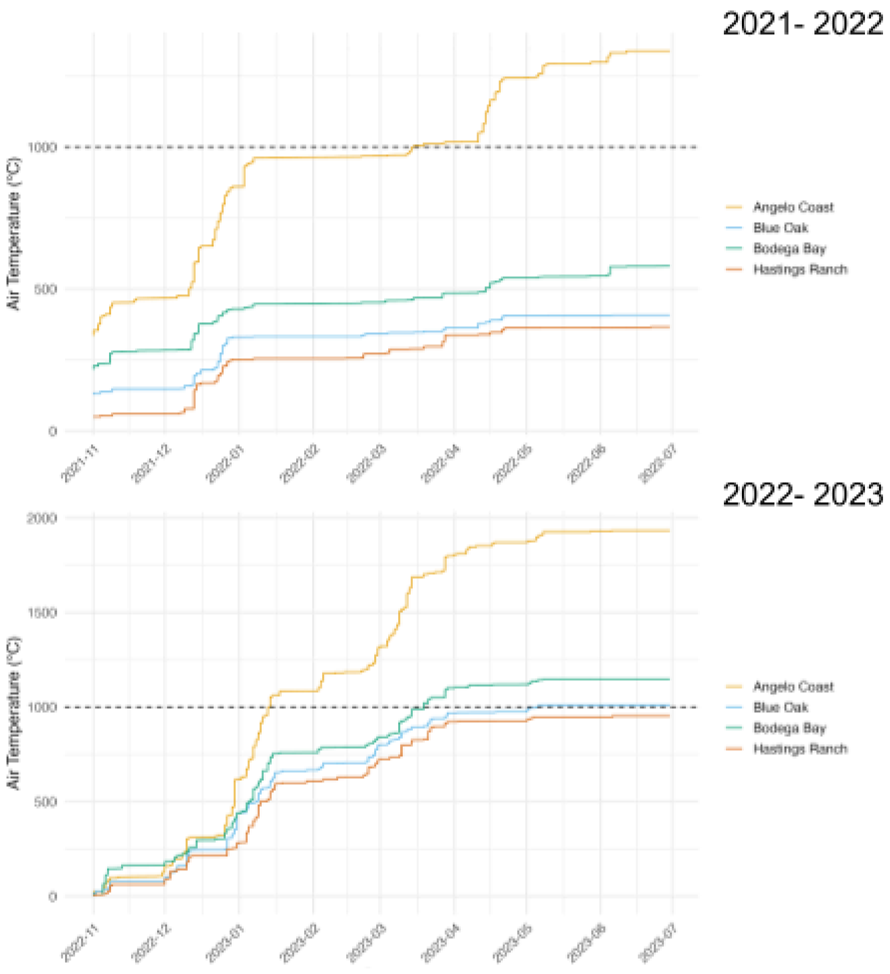


**Figure S6.** Cumulative rainfall (mm) at each of the four field sites.

Data is collected from mid-December to June for 2021-2022 (above) and 2022-2023 (below) from permanent sensors at each of the UC Reserves. The red line represents 1,000 mm on each graph to aid in comparison. Note that cumulative rainfall is higher in 2022-2023 compared to 2021-2022. AC receives significantly higher annual PPT (mm) than BB.

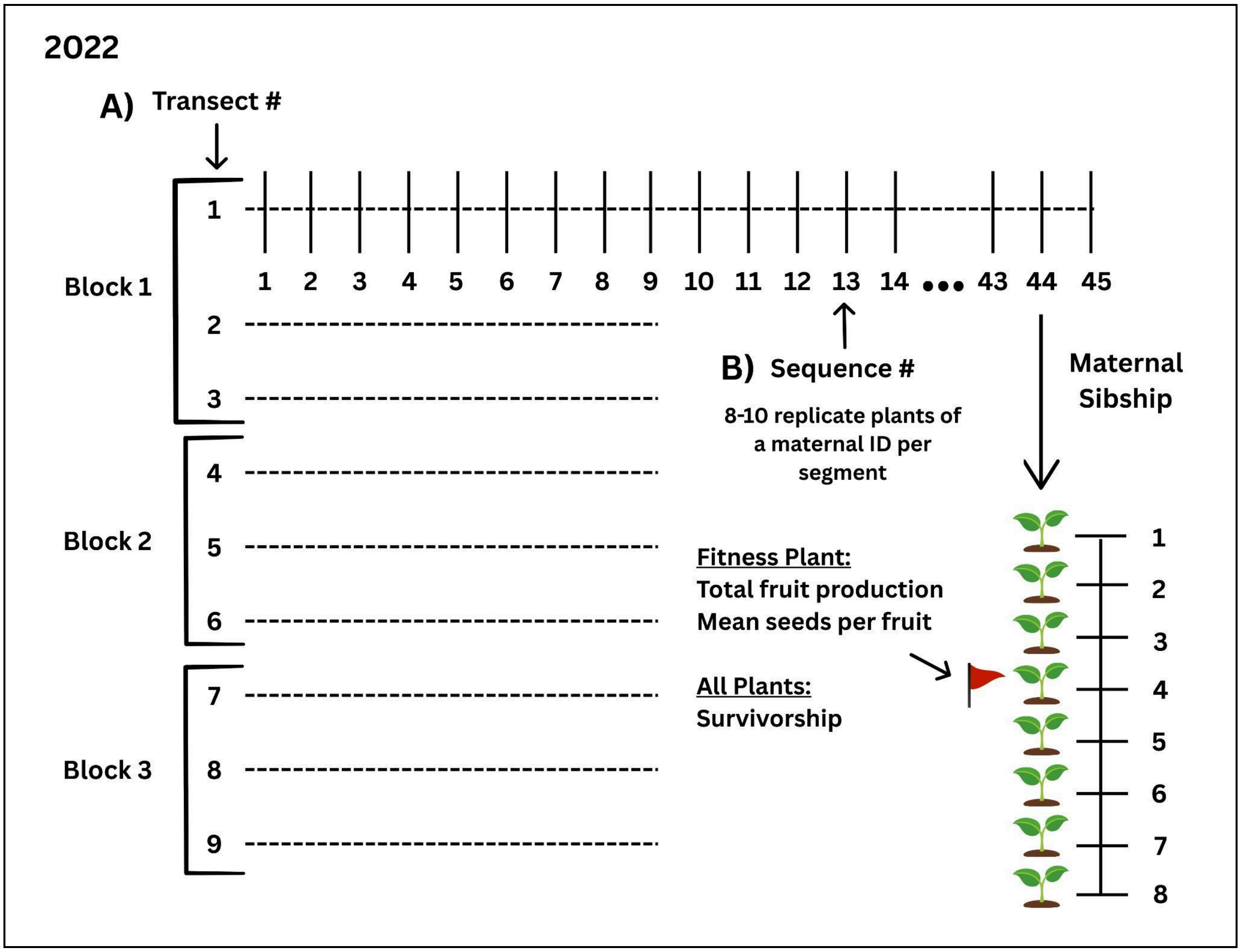


**Figure S7.** Experimental design at field sites

A) We had three blocks at each experimental site. Within each block, 3-4 transects (~45 meters long) were established, with planting segments spaced at 1-meter intervals along each transect. B) In Fall of year 1, 8-10 seeds from one maternal plant were sown at each segment, with each maternal sibship represented by one 1-meter segment in each block. When seeds were sown in Fall year 2, each segment contained seeds from two randomly selected maternal plants, with five seeds per maternal plant, drawn from the full set of generation 1 and generation 2 maternal families. In Fall year 2, generation 1 and generation 2 maternal families were assigned at random to all segments (block effects were not significant in year 1, and as a result, they were subsequently excluded in year 2), so each block did not necessarily contain the seeds of every maternal sibship. To distinguish between generations and sibships, we used labeled skewers and planted five seeds of a generation on different sides of the transect. As plants flowered in the winter and spring of year 1 and year 2, one plant per segment × maternal family × generation was randomly designated as the primary "fitness" plant, from which fitness-related trait data were recorded. Estimated sire effects on $W$ of parental family groups sown in 2022 and 2023 for the field sites AC, BB, BO, and HR.

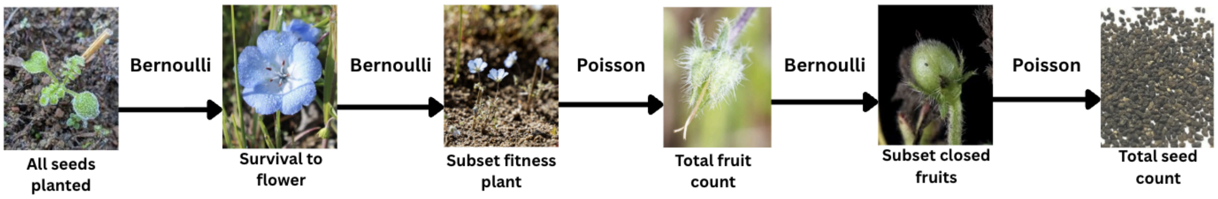


**Figure S8.** Aster graphical model used to estimate $\overline{W}$ for each generation at each site.

Arrows between life history stages indicate the statistical distributions conditional on the prior stage. The root node (all seeds planted) represents the 8-10 seeds sown per maternal plant at a segment. Survival to flowering and subsampling one fitness plant were modeled as Bernoulli distributions. Total fruit production and total seed count of intact fruits were assigned Poisson distributions, and the number of intact fruits (closed) was modeled as a Bernoulli distribution.

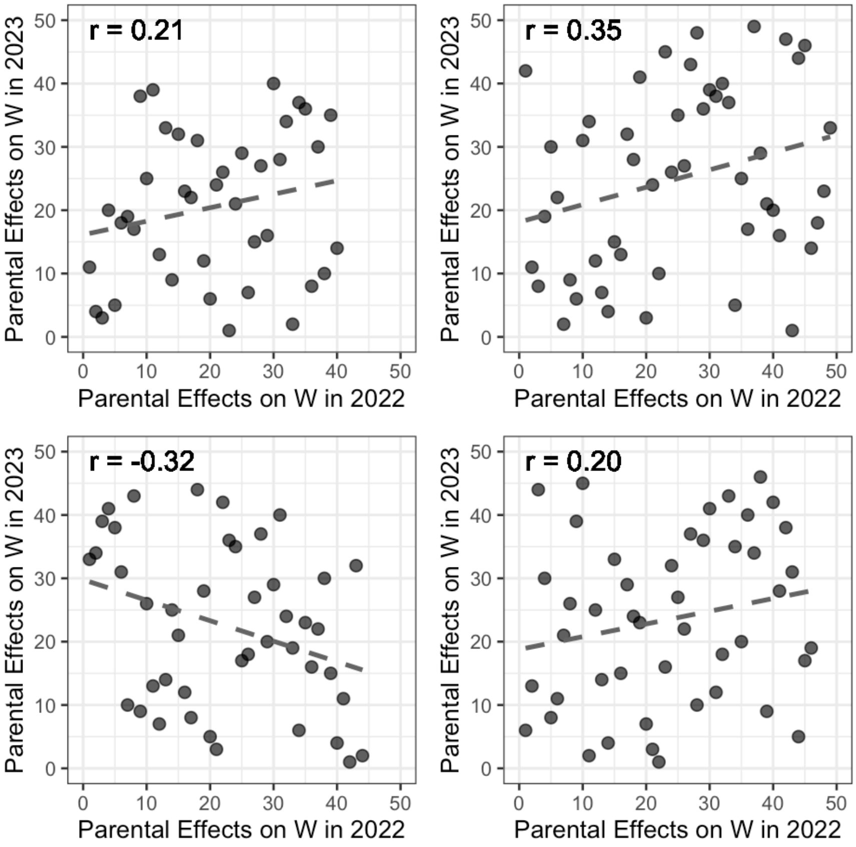


**Figure S9.** Bivariate Spearman rank correlation for estimated sire effects on $W$ of parental family groups sown in 2022 and 2023 for the field sites AC, BB, BO, and HR.

Bivariate Spearman rank correlation (and its correlation coefficient) among the sire effects on $W$ of the parental family groups in 2022 and 2023. These correlations do not account for variation within parental genotypes. Correlation coefficients: AC $r_s$ = 0.21, $p$ = 0.18; BB $r_s$= 0.35, $p$ = 0.01; BO $r_s$ = -0.32, $p$ = 0.03; HR $r_s$ = 0.20, $p$ = 0.18.

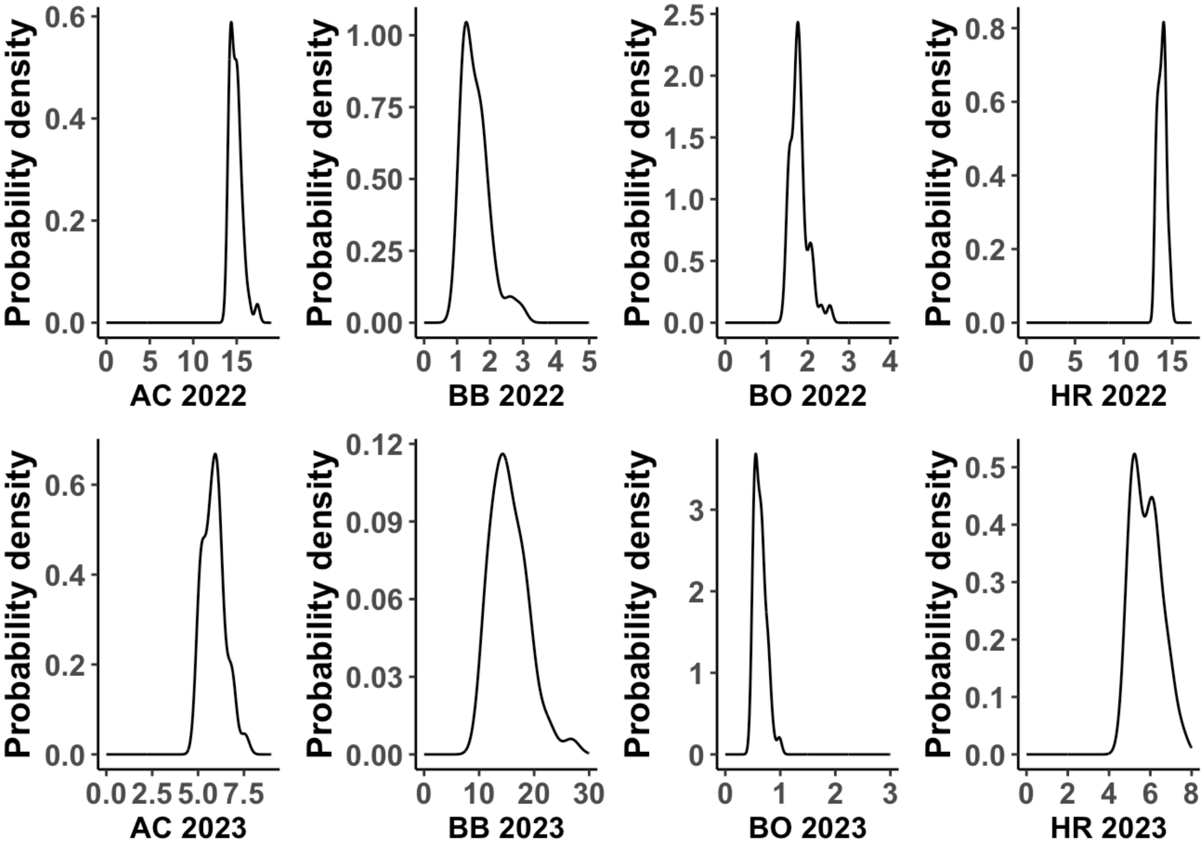


**Figure S10.** Probability densities of sire effects on fitness for the estimated mean lifetime fitness of parental sibships between years.

Each graph represents sire effects on fitness in generation 1 in both 2022 and 2023 population (AC = Angelo Cost; BB = Bodega Bay; BO = Blue Oak; HR = Hastings Reserve). The y-axis represents lifetime fitness estimates, and the x-axis represents probability densities. Note the scale for lifetime fitness for BB differs in 2023 due to high $\overline{W}$.

**Supplementary Tables:**

**Table S2.** Summary of sibships, transects, and seeds planted in generation 1.

Number of paternal and maternal sibships, transect number, segments per transect, plants per segment, and total seed number planted in Fall 2021 for each of the four populations (AC, BB, BO, HR) in generation 1.

| Population: | Paternal sibships: | Maternal sibships: | Transect / point #: | Segment # per transect/ | Seeds # per segment: | Total seed #: |
|---|---|---|---|---|---|---|

| | | | | point: | | |
|---|---|---|---|---|---|---|
| AC | 40 | 107 | 9 | ~36 | 10 | 3,210 |
| BB | 49 | 148 | 156* | ~3* | 8 | 3,528 |
| BO | 44 | 132 | 12 | ~44-23 | 8 | 3,168 |
| HR | 46 | 137 | 9 | ~46 | 8 | 3,288 |

**Table S3.** Population structure and seed planting overview for Fall 2022.

Number of paternal and maternal sibships, number from each generation (generation 1 and generation 2), transect number, segments per transect, plants per segment, and total seed number planted in Fall 2022 for each of the four populations (AC, BB, BO, HR).

| Population: | Paternal sibships: | Maternal sibships: | generation 1 seeds: | generation 2 seeds: | Transect/point #: | Segment # per transect/point: | Seeds # per segment: | Total seed #: |
|---|---|---|---|---|---|---|---|---|
| AC | 40 | 107 | 1,605 | 1,590 | 9 | ~36 | 5 | 3,195 |
| BB | 49 | 146 | 2,200 | 1,005 | 108* | ~2-4* | 5 | 2,205 |
| BO | 44 | 132 | 1,980 | 1,110 | 12 | ~44-23 | 5 | 3,090 |
| HR | 46 | 137 | 2,055 | 2,000 | 9 | ~46 | 5 | 4,055 |

**Table S4.** Parameter estimates from generation 1 and 2 aster fitness models.

Estimates from aster-based fitness models for generation 1 and generation 2 across four populations (AC, BB, BO, HR) in 2022 and 2023. The parental model includes effects of transect/location and parental variance on fitness, while the generation 2 model includes only the transect/location effect. Values represent model coefficients for each site-year combination.

| Generation 1 model | | | | Generation 2 model |
|---|---|---|---|---|
| Fitness ~ intercept + parental variance + transect | | | | Fitness ~ intercept + transect |
| **site** | **year** | **transect/ location** | **parental variance** | **transect/location** |
| AC | 1 (2022) | -0.002715 | 0.016874 | — |
| AC | 2 (2023) | -0.017395 | 0.039398 | -0.019521 |
| BB | 1 (2022) | -0.0011802 | 0.11509 | — |
| BB | 2 (2023) | -0.0009780 | 0.027294 | -0.0006052 |
| BO | 1 (2022) | 0.001068 | 0.11865 | — |
| BO | 2 (2023) | 0.015580 | 0.07184 | 0.023586 |
| HR | 1 (2022) | 0.004422 | 0.022499 | — |
| HR | 2 (2023) | 0.0004041 | 0.058031 | -0.0053172 |

**Table S5.** Likelihood ratio tests evaluating genotype-by-year (G × Y) interactions for fitness within each population.

G × Y interactions were assessed by comparing aster models with and without a parental × year random effect for generation 1 individuals within each population (AC: Angelo Coast, BB: Bodega Bay, BO: Blue Oak). ΔDeviance values, degrees of freedom (df), and associated p-values are reported for each population.

| Population | ΔDeviance | df | p-value |
|---|---|---|---|
| AC | 6.86 | 1 | 0.002 |
| BB | 47.56 | 1 | <0.001 |
| BO | 48.92 | 1 | <0.001 |

**Table S6.** Seed size and survival to flowering.

Logistic mixed models were fitted separately for each population, pooling both generations: survival to flowering ~ sown seed mass (mg) + generation + (1 | maternal line), using binomial errors. Odds ratios are reported per +1 mg increase in sown seed mass. Sown seed mass represents the $G_0$-weighed seed mass of the cross for generation-1 plants and the mean seed mass produced by the corresponding generation-1 family for generation-2 plants.

| Population | Plants (n) | Flowered (n) | Maternal lines | Slope (log-odds mg$^{-1}$) | SE | Odds ratio (95% CI) | Maternal-line SD | *P* |
|---|---|---|---|---|---|---|---|---|
| Angelo (AC) | 8,142 | 5,218 | 107 | 0.577 | 0.050 | 1.781 (1.616–1.963) | 0.515 | $2.9 \times 10^{-31}$ |
| Blue Oak (BB) | 6,781 | 2,600 | 144 | 0.022 | 0.017 | 1.022 (0.989–1.056) | 0.305 | 0.192 |
| Bodega Bay (BO) | 6,444 | 1,640 | 132 | 0.035 | 0.018 | 1.036 (0.999–1.074) | 0.322 | 0.055 |
| Hastings (HR) | 7,435 | 5,533 | 137 | 0.053 | 0.023 | 1.054 (1.007–1.104) | 0.460 | 0.023 |

**Table S7.** Mean seed size by generation.

Mean individual seed mass produced by plants in the generation-1 and generation-2 cohorts. The generation-1 cohort was sown from the original $G_0$ seed stock, whereas the generation-2 cohort was sown from seed produced by generation-1 plants during the 2022 field season. Both cohorts were then grown side by side during the 2023 field season, so generation is not confounded with year. Linear mixed models were fitted separately for each population: seed mass (mg) ~ generation + (1 | maternal line), with degrees of freedom and P-values estimated using the Satterthwaite approximation. The pooled model included site as an additional random effect: seed mass ~ generation + (1 | site) + (1 | maternal line). Thus, the estimated generation difference is adjusted for site, whereas the reported means are the raw values pooled across populations. Sample sizes include only plants with measurable seed mass (i.e., plants that produced seed), which is why they are smaller than the totals reported in Table S6.

| Population | G1 (n) | G2 (n) | G1 mean ± SD (mg) | G2 mean ± SD (mg) | Difference (G2 − G1) (95% CI) | df | *t* | *P* |
|---|---|---|---|---|---|---|---|---|

| | | | | | | | | |
|---|---|---|---|---|---|---|---|---|
| Angelo (AC) | 175 | 183 | 1.843 ± 0.788 | 1.748 ± 0.764 | −0.087 (−0.243 to 0.069) | 313.6 | −1.095 | 0.274 |
| Blue Oak (BB) | 238 | 108 | 3.616 ± 1.731 | 3.809 ± 1.894 | 0.269 (−0.125 to 0.663) | 317.1 | 1.340 | 0.181 |
| Bodega Bay (BO) | 104 | 41 | 3.772 ± 1.658 | 3.481 ± 1.664 | −0.291 (−0.890 to 0.309) | 136.8 | −0.950 | 0.344 |
| Hastings (HR) | 251 | 253 | 2.335 ± 1.122 | 2.285 ± 1.633 | −0.063 (−0.301 to 0.175) | 440.8 | −0.517 | 0.605 |
| **All populations (pooled)** | **768** | **585** | **2.815 ± 1.569** | **2.482 ± 1.662** | **−0.023 (−0.174 to 0.127)** | **1177.8** | **−0.302** | **0.763** |
